# Structure and quenching of a bundle-shaped phycobilisome

**DOI:** 10.1101/2025.06.10.658930

**Authors:** Anna D. Burtseva, Yury B. Slonimskiy, Timur N. Baymukhametov, Maria A. Sinetova, Daniil A. Gvozdev, Georgy V. Tsoraev, Dmitry A. Cherepanov, Eugene G. Maksimov, Vladimir O. Popov, Konstantin M. Boyko, Nikolai N. Sluchanko

## Abstract

Cyanobacteria use soluble antenna megacomplexes, phycobilisomes (PBS), to maximize light-harvesting efficiency and small photoswitchable orange carotenoid proteins (OCPs) to down-regulate PBS in high light. Among known PBS morphologies, the one from the basal cyanobacterial genus *Gloeobacter* still lacks detailed structural characterization. Here, we present the cryo-EM structure of the *G. violaceus* PBS, a >10-MDa complex with a unique architecture consisting in 516 polypeptide chains totaling nearly 90,000 residues and harboring over 860 bilin chromophores. This unique PBS features diverging, conformationally mobile bundles of rods composed of stacked phycoerythrin and phycocyanin hexamers, stemming from a pentacylindrical allophycocyanin core belted by auxiliary phycocyanin hexamers. We show how two *Gloeobacter*-specific multidomain linker proteins, Glr1262 and Glr2806, maintain this bundle-shaped architecture and reveal its differential regulation via non-photochemical quenching by two OCP types of *G. violaceus* that recognize separate binding sites within the allophycocyanin core, including lateral cylinders absent in tricylindrical cores. Our findings provide the high-resolution structural insight into *Gloeobacter* PBS and its regulation, revealing divergent adaptations in early-branching cyanobacteria. The structure advances understanding of PBS diversity and evolution, offering a framework for bioengineering light-harvesting systems in synthetic biology and biotechnological applications.

---

Light absorption is the first step in photosynthesis, and photosynthetic organisms have evolved various mechanisms to improve its efficiency. Cyanobacteria and red algae use extramembranous antenna megacomplexes called phycobilisomes (PBS), composed of phycobiliproteins (PBPs), to harvest light. PBPs covalently bind bilins, which are open-chain tetrapyrroles (*1*). Bilins absorb light in different spectral regions and transfer energy to the reaction centers of photosystems (*2–4*). This compensates for chlorophyll’s limited ability to absorb light at short wavelengths (*5*). The efficient funneling of energy absorbed by bilins in PBPs is enabled by structural organization of PBS. PBPs are composed of heterodimeric αβ protomers, often designated “monomers”, which oligomerize into ring-like trimers (αβ)_3_ and hexamers (αβ)_6_. These further assemble with the help of linker proteins that interact with the PBP-rings in their cavities. A typical PBS contains an allophycocyanin (AP) core – two (*6*), three (*7*), or five (*8*) cylinders of (αβ)_3_ trimers of AP – and a set of radiating rods – stacks of hexamers formed by αβ protomers of phycocyanin (PC) and, in red algae and some cyanobacteria, phycoerythrin (PE) (*1*, *9*, *10*). AP, PC, and PE differ in the number of bilins bound to their α (one in AP and PC, or two in PE) and β (one in AP, two in PC, and three in PE) subunits. The three main bilin types in cyanobacterial PBS are phycocyanobilin (PCB, found in AP and PC), phycoerythrobilin (PEB), and phycourobilin (PUB) (both found in PE) (*1*), whose collective absorbance covers the visible spectral range that is poorly accessible to chlorophyll.

Known PBS in red algae are hemi-ellipsoidal/block-type (*10*, *11*), whereas in cyanobacteria the prevalent PBS form is hemi-discoidal (*1*). In addition, the paddle-shaped PBS was recently described in the relict cyanobacterium *Anthocerotibacter panamensis* UTEX 3164, which utilizes single PC hexamers and chains of staggered PC hexamers that surround a heptacylindrical AP core instead of rods (*5*). A unique case is the cyanobacterium *Gloeobacter violaceus* PCC 7421 (hereafter, *G. violaceus*), which diverged from the crown cyanobacteria the earliest (∼2 billion years ago). *G. violaceus* is characterized by slow metabolism and growth; primitive bacterial-type carotenoid biosynthesis; an inability to stably maintain intracellular pH; and a lack of circadian clock and thylakoids (*12*, *13*). PBS in *G. violaceus* attach to the photosystems localized in the plasma membrane (*12*, *14*). Although bundle-type PBS were first described in 1981 (*12*), they remain the only known PBS morphology without a high-resolution structure. The regulation of the bundle-shaped PBS in high light also remains unknown.

## 1. Cryo-electron microscopy of the bundle-shaped *G. violaceus* PBS

The *G. violaceus* culture exhibited an absorbance spectrum with peaks at 500, 560, and 620 nm, corresponding to PBPs. Small features were also present at 440 and 680 nm, indicating relatively low chlorophyll content (Fig. 1A) (*14*). *G. violaceus* PBS (hereafter, GviPBS), which was isolated via sucrose gradient ultracentrifugation (Fig. 1B), exhibited the same prominent absorbance bands as PBPs, reflecting the presence of PUB (500 nm), PEB (560 nm), and PCB (an excitonic peak at 650 nm and a shoulder at 620 nm) (Fig. 1A). The first two of these are more prevalent in red algae than in cyanobacteria (*1*, *11*). All resolved cyanobacterial PBS structures contain only PCB and no other bilin types (*1*). PBPs could be biochemically separated into three distinct PBP fractions, corresponding to PE, PC, and AP, according to their absorbance and fluorescence spectra (Fig. 1C). The presence of PBPs and all linker proteins, including *Gloeobacter*-specific multidomain linkers Glr1262 and Glr2806, was confirmed by SDS-PAGE and mass spectrometry (Fig. S1 and Table S1).

**Fig. 1.**
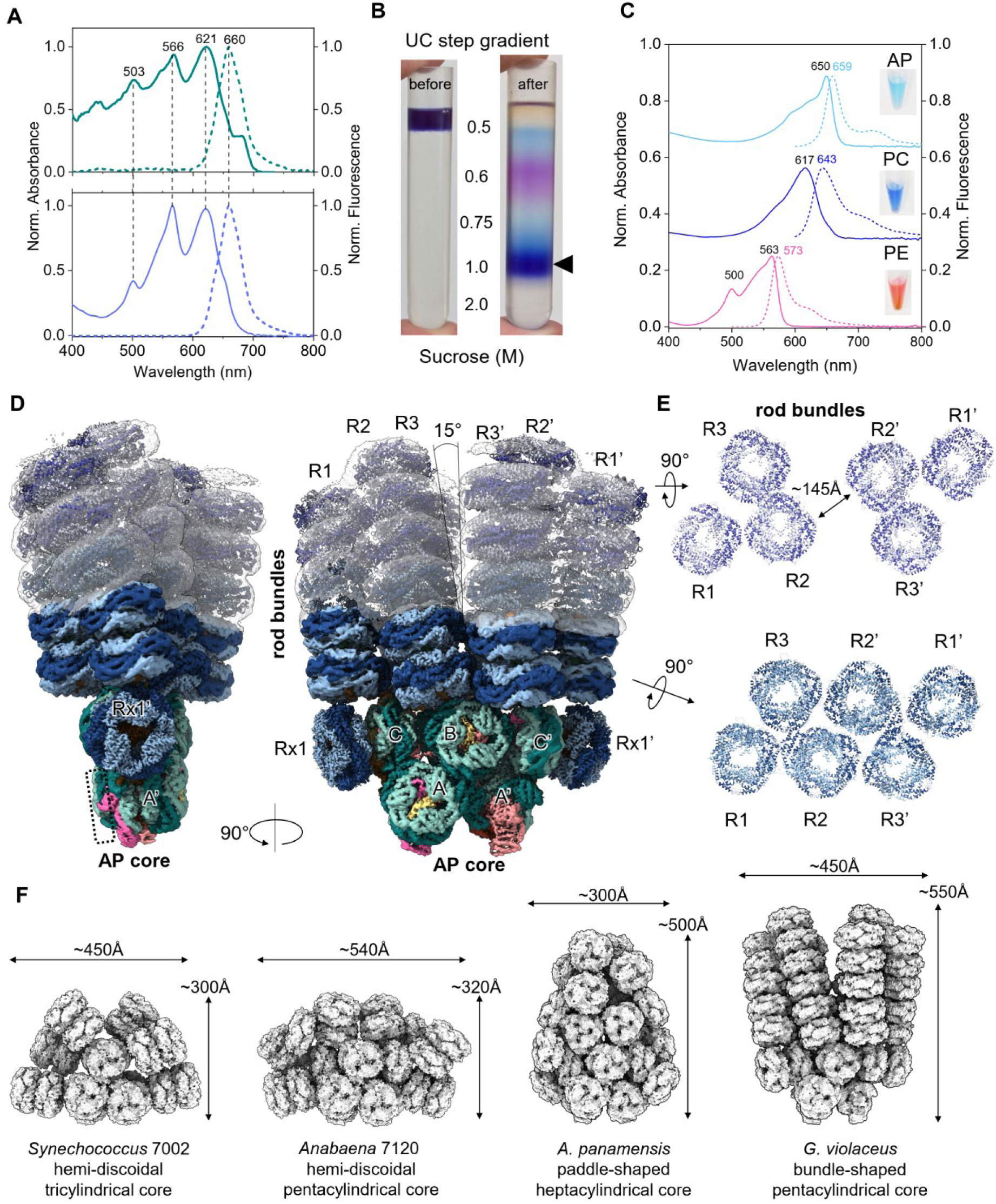
The bundle-shaped phycobilisome from *G. violaceus*. **A**. Absorption (solid line) and fluorescence (dashed line) spectra of *Gloeobacter* cells (top) and isolated PBS (bottom). An integrating sphere was used to measure the absorption spectrum of the cells. In both cases, fluorescence was excited at 630 nm. **B**. PBS isolation by ultracentrifugation in sucrose gradient as shown. Arrow indicates the PBS zone. **C**. Absorption (solid line) and fluorescence (dashed line) spectra of the individual allophycocyanin (AP), phycocyanin (PC) and phycoerythrin (PE). Excitation was at 475 (PE) or 575 nm (PC, AP). The samples are shown in insert. **D**. Composite cryo-EM map of the bundle-shaped PBS showing the location of the AP core (A/A’, B and C/C’ cylinders) and rod bundles. **E**. Sections across the rods on the two levels indicated by arrows, showing diverging three-rod bundles R1-R2-R3 and R1’-R2’-R3’. **F**. Cryo-EM structures representing the relative sizes of different cyanobacterial PBS – hemi-discoidal PBS from *Synechococcus* and *Anabaena* (*8*), paddle-shaped PBS from *A. panamensis* (*5*) and bundle-shaped PBS from *G. violaceus* PBS (this work).

To preserve the intact PBS for cryo-EM, we performed ultracentrifugation in parallel gradients of sucrose and glutaraldehyde (GraFix (*15*)) (Fig. S1). To improve the quality of the cryo-EM density map, we focused on a set of substructures for model building and refinement against the masked map. This resulted in six focused maps with resolutions ranging from 2.72 Å to 3.67 Å (Table S2, Fig. S2, S3). The refined substructures were then combined to create a model of the entire PBS and fitted into the composite cryo-EM map (Fig. 1D).

For the GviPBS model, we used the crystallographic structures of (αβ)_6_ hexamers of AP (PDB 2vjt), PC (PDB 2vjr) and PE (PDB 2vjh) (Fig. S4) and available electron microscopy and mutagenesis data (*16*). The reconstructed cryo-EM structure revealed three pairs of C2-symmetry-related rods stemming from the top of the pentacylindrical AP core (R1/R1’, R2/R2’, R3/R3’) (Fig. 1D,E). The AP core contains two antiparallel bottom cylinders (A/A’), a top cylinder (B) and two lateral AP (αβ)_6_ hexamers (C/C’), which are attached sideways to the top B cylinder. This “3+2” AP core can be superimposed on the pentacylindrical cores in hemi-discoidal PBS from *Anabaena* sp. PCC 7120 (*8*) and *Thermosynechococcus vulcanus* NIES-2134 (*17*) as well as on the lower portion of the heptacylindrical AP core in *A. panamensis* PBS (*5*) (Fig. S5). In GviPBS, the “3+2” core extends to the sides with additional PC (αβ)_6_ hexamers (Rx1/Rx1’) that lie coaxially with the C/C’ (αβ)_6_ hexamers of AP and support the rods (Fig. 1D).

The cryo-EM map of the rods dispersed more towards the periphery. Prior work revealed that deleting PE subunits results in a shorter GviPBS morphology, with rods containing three PC (αβ)_6_ hexamers in height (*16*). Although two PC (αβ)_6_ hexamer stacks could be confidently built in each rod, the density of the third PC hexamer and PE (αβ)_6_ hexamers only permitted approximate placement (Fig. 1D). Two (αβ)_6_ hexamers of PE (PDB 2vjh) could be docked in the rods R1/R1’ and R3/R3’, whereas three PE (αβ)_6_ hexamers could be docked in the rods R2/R2’, for a total of 18 PC and 14 PE (αβ)_6_ hexamers in the rods of the composite model. Unexpectedly, the rods formed two bundles (R1-R2-R3 and R1’-R2’-R3’) that radiated at ∼15° angle so that the first-level PE (αβ)_6_ hexamers in rods R2/R2’ became ∼145 Å apart (Fig. 1D,E). Each rod bundle formed a superhelix with a period of ∼1000 Å (Fig. S6). Two maps supporting different bundle conformations allow us to imagine movement of the bundles relative to each other (Fig. S7 and Movie S1).

The cryo-EM map permitted confident modeling of seven AP (αβ)_6_ hexamers out of eight, while the fourth (αβ)_3_ trimer was absent from both bottom AP cylinders (Fig. 1D), similar to *T. vulcanus* PBS (PDB 7vea) (*17*). The absence of the fourth (αβ)_3_ trimer may be due to its loss during sample preparation (*1*, *17*, *18*), though light-controlled reconfiguration of the PBS architecture is also possible (*19*). Notably, the missing trimer would contain ApcD, one of the terminal emitters responsible for the energy transfer to the photosystems in other PBS (*6–8*). Mass spectrometry confirmed that both minor PBPs, ApcD (α-chain) and ApcF (β-chain), were present in our GviPBS preparations (Table S1). Since no justification based on density alone was found for an alternative location of ApcD (instead of the common location in the fourth trimer), ApcD was not included in our models. ApcF was modeled in its expected position in the third AP trimer of the basal cylinders (*6–8*), although the corresponding density did not allow us to distinguish it unambiguously from ApcB because the two proteins are highly identical (70.8%). Notably, *A. panamensis* PBS naturally lacks ApcD and ApcF (*5*). Similarly, deleting either gene does not affect the functionality or regulation of PBS in *Synechocystis* sp. PCC 6803 (hereafter *Synechocystis* 6803) (*20*). Thus, we assume that the fourth ApcD-containing AP trimer is dispensable for GviPBS functioning.

The AP core width is narrower than the lower level of the rods (Fig. 1D). However, blurred density proximal to Rx1/Rx1’ suggests the tentative placement of Rx2/Rx2’ (Fig. S8). Consistent with the mutagenesis data (*16*), we hypothesize that there are up to three Rx hexamers on each side of the core. The third Rx substructure is likely too dynamic to have a distinct density (Fig. S8). The tentative positioning of three Rx hexamers is sterically compatible with the lateral stacking of GviPBS into arrays, similar to those of PBS in crown cyanobacteria (*3*, *7*) (Fig. S8). This scenario is most probable in thylakoid-less *G. violaceus* because it saves space on the plasmalemma, which accommodates all photosynthetic complexes in this organism (*12*, *14*).

## 2. Linker proteins

The reconstructed GviPBS structure contains 82 (αβ)_3_ trimers of PBP and 864 bilins. It is the largest resolved cyanobacterial PBS, with dimensions of 450×550 Å, exceeding those of paddle-like and hemi-discoidal PBS (Fig. 1E). The bundle-shaped morphology is maintained by linker proteins (Fig. 2A), including the multidomain linkers Glr1262 and Glr2806, which are unique to *Gloeobacter* (*16*, *21*, *22*). Each is predicted to have three pfam00427 (REP) domains in its sequence (*16*, *21*). Nevertheless, these proteins are only 58.4% identical to each other and are dissimilar to the functional analogs CpcG and CpcL from *Synechocystis* 6803 (<30% sequence identity). We identified the precise location of Glr1262 and Glr2806 REP domains by cryo-EM density for non-conserved amino acids (Fig. S9, S10). Glr2806 is responsible for attaching Rx to the core (*16*). Its long N-terminal arm interacts with the A/A’ and C/C’ cylinders, while its REP1 domain binds inside the Rx1/Rx1’ cavity (Fig. 2A-C). The REP2-REP3 domains of Glr2806 are expected to tether two additional Rx copies, Rx2/Rx2’ and Rx3/Rx3’, to the core (Fig. S8) (*16*). Although mass spectrometry could not cover the entire Glr2806 sequence (Table S1), our data unequivocally confirm Glr2806’s role in attaching Rx structures to the core (Fig. 2B). Interestingly, Glr2806 of *G. morelensis* MG652769 also contains three REP domains, but the *G. kilaueensis* JS1 homolog lacks one (Fig. S11). This suggests that three Rx hexamers are not necessary for bundle-shaped PBS assembly.

**Fig. 2.**
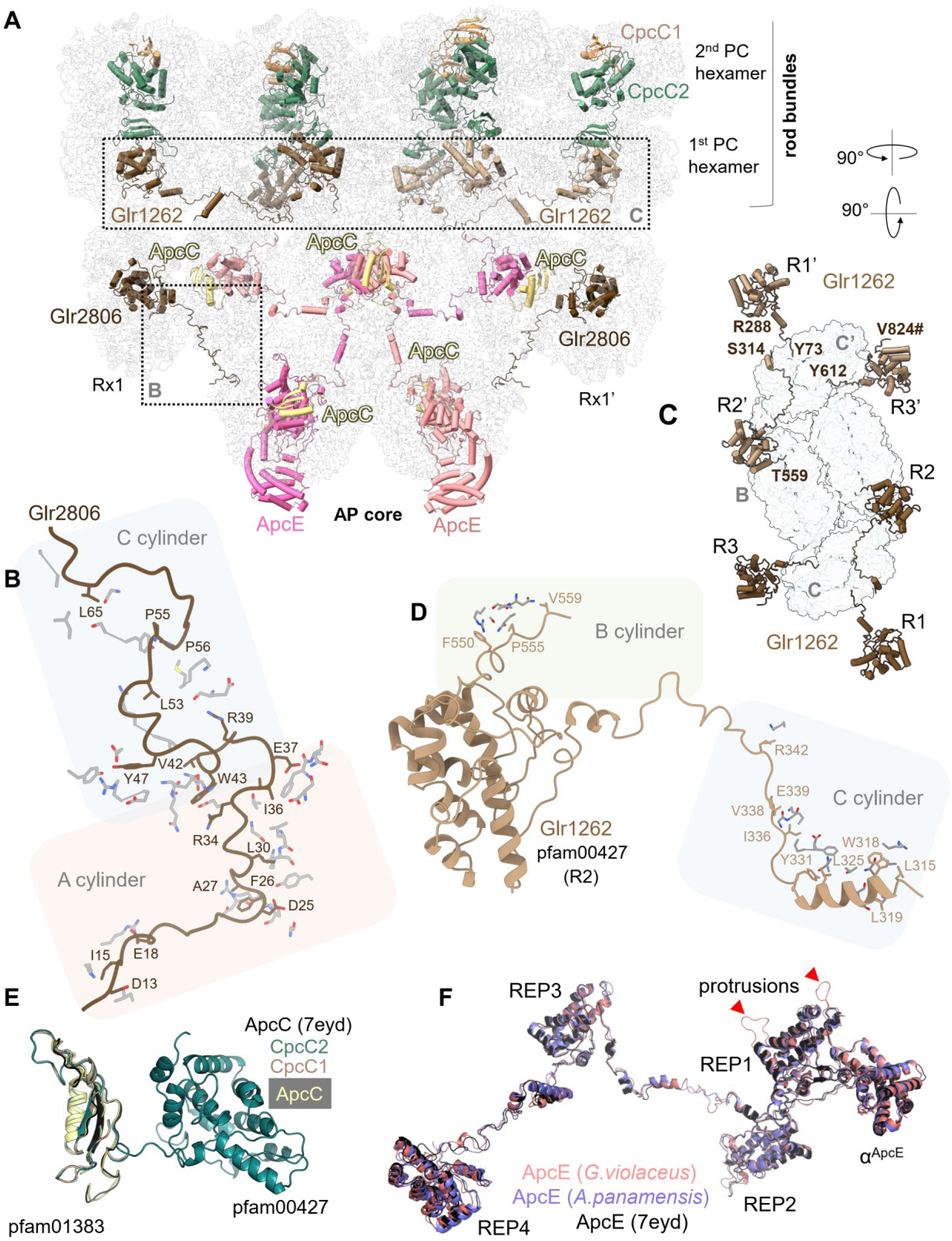
Linker proteins of GviPBS. **A**. Front view of the AP core and two PC levels in the rods. PBP hexamers are shown as semitransparent ribbons, linker proteins are in cartoon representation. All linkers are labeled except for CpcC1/CpcC2 for which only one copy is labeled. Parts of the model for focused analysis in separate panels are framed. **B**. Contacts formed by the N-terminal arm of Glr2806 with the core cylinders A/A’ and C/C’. **C**. Topology of the two copies of Glr1262 on top of the core showing relative positions of the repeat domains REP1, REP2, REP3 that attach bases of the rods R1/R1’, R2/R2’, R3/R3’, respectively, by clustering them as two rod bundles. # indicates the C terminal of the Glr1262 chain. **D**. Contacts made by the arms of Glr1262 with the core cylinders B and C/C’. **E**. Superposition of ApcC capping linker of *G. violaceus* with ApcC from Anabaena PBS (PDB 7eyd) and with the equivalent pfam001383 domains of CpcC1 and CpcC2 rod linkers. **F**. Superposition of ApcE from *G. violaceus*, *A. panamensis* and *Anabaena* 7120 showing the location of the REP1-4 domains and the bilin-binding α^ApcE^ domain. The profound difference is the presence of large unstructured protrusions exclusively in REP1 of *G. violaceus* ApcE (indicated by red arrows).

The Glr1262 linker is also predicted to have three pfam00427 domains, which enable the interaction with three PBP hexamers. Each REP domain of Glr1262 is superimposable with Glr2806 REP1 (0.6-1 Å Cα RMSD; Fig. S12). The density helped unequivocally localize Glr1262 on the first level of PC hexamers in the rods, consistent with its predicted position (*16*). The three REP domains (pfam00427) of Glr1262 were identified at vertices of an equilateral triangle, which corresponded to the cavities of three PC hexamers that formed the base of a rod bundle (Fig. 2C). Although incomplete density prevented tracing the entire Glr1262 chain, assigning REP domains to a Glr1262 copy that connects neighboring PC hexamers within the rod bundle appears to be the most reasonable option. Two copies of Glr1262, each with three REP domains, are necessary and sufficient to cluster the six rods symmetrically into two triangular rod bundles. Each triangle is centered at the top side of the C/C’ cylinder of AP (Fig. 2C). In the absence of a CpcG homolog in the genome, a rod-core linker found in studied cyanobacteria, Glr1262 apparently plays this role in *Gloeobacter* (*16*, *21*). Indeed, the REP1-REP2 arm (residues 280-350) of Glr1262 interacts with the C/C’ cylinder residues and the REP2-REP3 arm interacts with the B cylinder of the AP core (Fig. 2D).

We located the conserved rod linkers, CpcC1 and CpcC2 (*22*), which contain pfam01383 and pfam00427 domains that connect two neighboring PC layers (Fig. 2A, E). According to mass spectrometry, all small capping linkers were present in the sample (Table S1). However, we could unequivocally localize multiple copies of only the core linker ApcC (Fig. 2A). Their pfam01383 domain is similar to those of other PBS types and to the pfam01383 domain in CpcC1/CpcC2 (Fig. 2E).

The pentacylindrical GviPBS core is assembled by two copies of the core-membrane linker ApcE (or L_CM_), which has four conserved REP domains (pfam00427) that bind within the cavities of four AP hexamers (Fig. 2F). Pfam00427 domains of ApcE are conformationally similar to REP domains of Glr2806, Glr1262 and CpcC1/CpcC2 (<1.6 Å Cα RMSD) (Fig. S12). Although ApcE of GviPBS is conformationally similar to other ApcE proteins from PBS with the pentacylindrical cores, it contains unique unstructured protrusions that potentially interfere with the attachment of the fourth ApcD-containing trimer (Fig. 2F). As in other known cyanobacterial PBS structures, ApcE is the only linker in GviPBS with a chromophore. It is canonically located within the N-terminal domain of ApcE, incorporated into the third AP (αβ)_3_ trimer of the basal cylinders as the α subunit.

## 3. Chromophore system

GviPBS is notable for containing not only PCB, but also PEB and PUC chromophores, which give *G. violaceus* its characteristic violet color. PCB is located in AP core and PC hexamers in rods, whereas PEB and PUC are solely located within the upper part of the rods (Fig. 1D). The PEB and PUC chromophores could not be sufficiently resolved in cryo-EM maps; therefore, their conformations and local environments were inferred from the crystal structure of PE (PDB 2vjh). PCB conformations within PC hexamers in rods and AP hexamers in the core were resolved in cryo-EM density map, including the terminal emitting bilin in ApcE (Fig. 3A-E). The three bilin types differ in the number of conjugated double bonds, increasing from PUB to PEB to PCB, which is accompanied by enhanced coplanarity of the pyrroles (Fig. 3F,G). While PCB and PEB are covalently attached to the single conserved cysteines of the α and β subunits, PUB is doubly tethered to two cysteines simultaneously (Fig. 3A-E). The deficit of double bonds between the pyrroles and the two-sided covalent linkage forces PUB to adopt a bent conformation, with A and D rings completely outside the B-C plane (Fig. 3A). This is consistent with the absorbance maximum of ∼500 nm for PUB, the shortest among *G. violaceus* PBPs (Fig. 1C) (*9*). PEB has two double bonds in the linkages between the pyrrole rings, with A and D rings displaced on the opposite sides of the B-C plane (Fig. 3B,F), featuring a red-shifted absorbance maximum of 560 nm. PCBs with the three double bonds in the linkages between the four pyrrole rings adopt conformations with different degrees of ring coplanarity and the optional presence of hydrophobic side chains in the vicinity of the D ring. The ApcE bilin conformation has the most coplanar A-B-C-D rings. Its A ring flips from a *ZZZasa* to a *ZZZssa* configuration (*23*), while the D ring remains coplanar due to a π-stacking interaction with the Trp162 side chain (Fig. 3E), missing from PCB-binding sites in other AP or PC hexamers (Fig. 3A-D). Correspondingly, the *ZZZssa* bilin of ApcE is a red-shifted fluorophore with the properties of the terminal emitter (*23*).

**Fig. 3.**
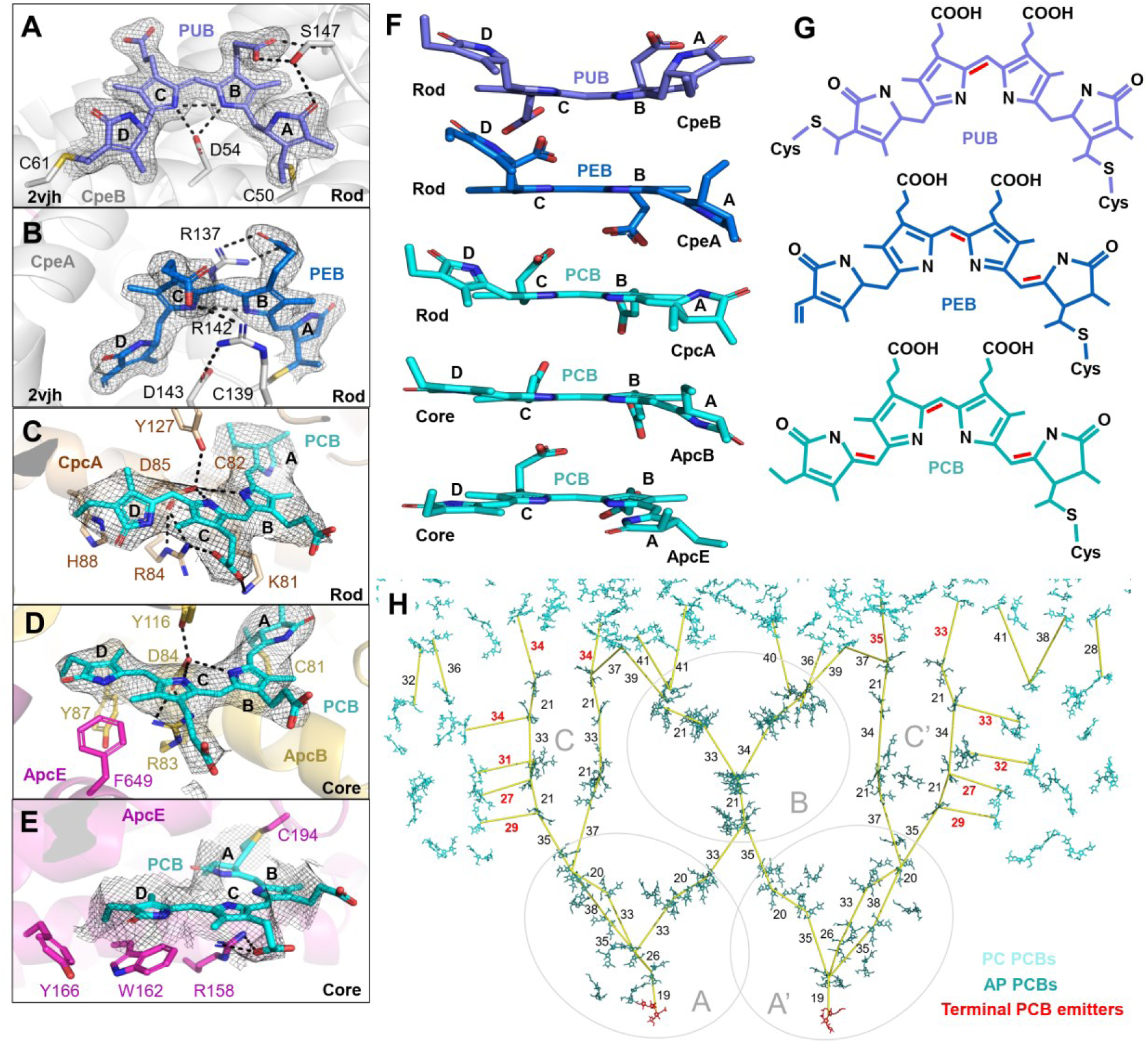
Bilins of GviPBS. **A-E.** Representative PUB, PEB and PCB bilin conformations, shown with the polar contacts to the surrounding protein residues and the respective electron density (1σ contour). PUB and PEB binding sites are imaged using the crystal structure of PE (PDB 2vjh), PCB sites are from cryo-EM data obtained in this work. **F**. Coplanarity of the bilins shown in panels **A-E**. Pyrrole rings A-B-C-D are labeled. **G**. Structures of the three bilin types found in GviPBS. Red lines mark the distinctive double bonds connecting the pyrroles. **H**. Estimation of the EET pathways from PC to AP and within AP in GviPBS. Distances connecting mass centers of the selected PCBs are indicated in Å by yellow lines. Remarkable shortest distances (≤35 Å) between PC and AP bilins are indicated by red font. Terminal emitting bilins of ApcE are shown by red sticks.

The stoichiometry of bilin incorporation (1:1 per αβ protomer of AP, 1:2 per αβ protomer of PC, 2:3 per αβ protomer of PE) establishes a descending bilin concentration gradient from the top to the button of GviPBS (Fig. S13), which may be relevant for intra-PBS excitation energy transfer (EET). EET pathways are typically estimated based on minimal interbilin distances (*5*, *6*, *8*). The bundles of rods position multiple PCB chromophores within 35 Å of each other, which is sufficient for interrod EET. This increases the probability of capturing photons in typical *G. violaceus* habitats (*1*, *14*, *16*). However, the absence of lateral radiating rods in *G. violaceus*, which enable direct EET to the A/A’ core cylinders in crown cyanobacteria (*7*, *8*), suggests peculiarities in rod-core EET in *G. violaceus*.

Remarkably, the minimal distances (∼33-35 Å) between the rod and core bilins in GviPBS were found to be between the PCBs of the lowest PC hexamers and the upper PCBs of the AP hexamers of C/C’ cylinders (Fig. 3H). The corresponding distances involving PCBs of the top B core cylinder are 37-41 Å. Furthermore, the coaxial position of PC hexamers Rx1/Rx1’ with C/C’ core cylinders brings several PCBs of those PC hexamers to distances of 27-31 Å from AP bilins (Fig. 3H). This is expected to substantially facilitate EET and indicates the special relevance of C/C’ hexamers (a hallmark feature of the pentacylindrical cores) in receiving energy from rods and auxiliary PC hexamers in GviPBS.

## 4. Non-photochemical quenching of the bundle-shaped PBS

*G. violaceus* grows optimally under low insolation but is known to respond to intense blue light by inducing non-photochemical quenching (NPQ) at the PBS level (*24*). High light threatens to damage the photosynthetic apparatus of cyanobacteria by increasing the risk of the reactive oxygen species formation. To protect themselves, cyanobacteria express a unique orange carotenoid protein (OCP). In high light, OCP converts from an inactive orange (O) into an active red (R) form that binds to the PBS core and dissipates the excess absorbed energy as heat (*7*, *25*). OCP is composed of an N-terminal domain (NTD) and a C-terminal domain (CTD) that share a non-covalently bound ketocarotenoid such as echinenone (ECH) (*26–30*). OCP-mediated photoprotection has been thoroughly studied in vivo (*25*, *31*, *32*) and reconstituted in vitro (*33*); however, the OCP-binding site has only been structurally elucidated for the hemi-discoidal PBS of *Synechocystis* 6803 (SynPBS) (*7*, *34*), which has a tricylindrical AP core (Fig. S14). Nevertheless, OCP-mediated quenching of the pentacylindrical PBS cores should differ, because C/C’ core cylinders completely obscure the binding site for one of the OCP subunits (Fig. S14).

There are three main OCP clades: OCP1, OCP2 and OCP3. The latter is subdivided into subclades a, b and c, with OCP3a being the closest to a presumed OCP ancestor (Fig. 4A) (*28*, *30*, *35*, *36*). The only characterized *Gloeobacter* OCP is found in *G. kilaueensis* JS1, which encodes only one OCP type: GkilOCP3a (*35*) (Fig. 4B). *G. morelensis* and *G. violaceus* contain two uncharacterized copies of the full-length OCP3a, which have only 66.6% sequence identity. These variants are designated here as OCP3a_α (apparently equivalent to GkilOCP3a) and OCP3a_β (Fig. S15).

**Fig. 4.**
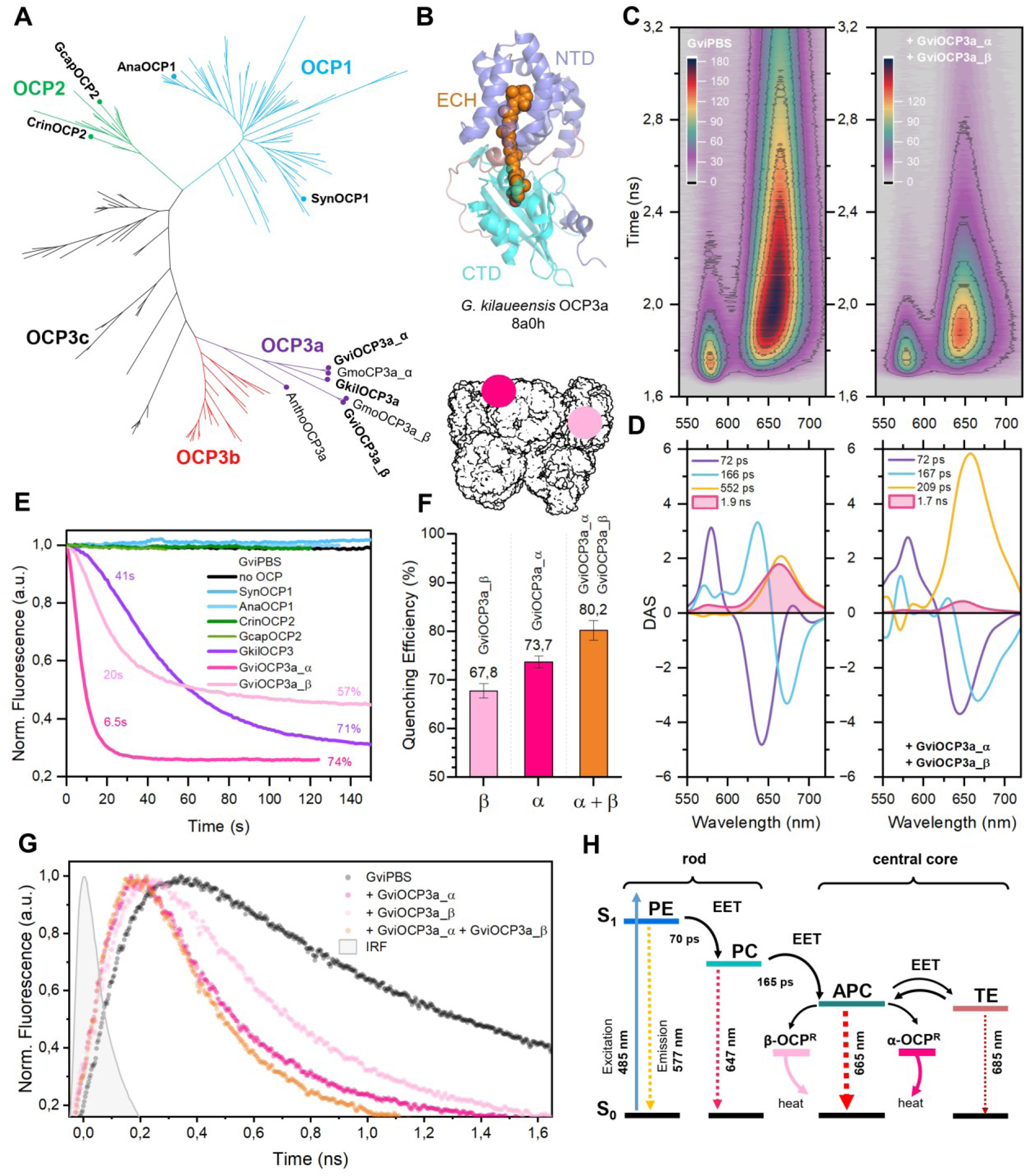
Quenching of GviPBS by orange carotenoid proteins. **A**. Phylogenetic tree showing the relationships between OCP1, OCP2, OCP3a, OCP3b and OCP3c clades. OCP homologs considered in this work are labeled. Construction of the tree is described in detail in Methods. **B**. Crystal structure of OCP3a from *G. kilaueensis* (*35*) showing the location of its NTD and CTD domains and the embedded echinenone (ECH) chromophore required for photoactivity. **C**. Time-resolved emission spectra (TRES) of GviPBS fluorescence in the absence and presence of the combination of GviOCP3a_α and GviOCP3a_β. **D**. Decay associated spectra (DAS) of GviPBS fluorescence data presented in **C**. The numbers indicate characteristic kinetic components derived from the global analysis of time-resolved fluorescence spectra (**C**) showing the response to photoactivation of GviOCP3a_α and GviOCP3a_β in combination. **E**. Time-courses of GviPBS quenching by OCP variants, showing characteristic time constant of 2-fold reduction of fluorescence and overall quenching efficiency. **F**. Quenching efficiency estimated as a relative reduction of yield of c.a. 1.8 ns components upon the photoactivation of GviOCP3a_α and GviOCP3a_β and their combination based on global analysis of TRES (C). **G**. Normalized fluorescence decay kinetics of GviPBS at 680 nm excited by 150 fs laser pulses at 485 nm. The temperature of the samples was stabilized at 10 °C during all experiments to prevent fluorescence recovery. **H**. Schematic description of the excitation energy transfer (EET) processes within GviPBS and the effect of GviOCP3a_α and GviOCP3a_β on dissipation of the absorbed energy.

We produced GviOCP3a_α and GviOCP3a_β proteins recombinantly and confirmed that, as expected for native OCP proteins in *Gloeobacter* (*35*), they contained exclusively ECH as the carotenoid. Both proteins were monomeric and exhibited similar absorbance spectra and photoactivity (Fig. S16). To compare PBS quenching by these proteins, we employed steady-state and picosecond time-resolved emission spectroscopy (Fig. 4C,D,E and Fig. S17). OCP1 from *Synechocystis* 6803 (SynOCP1) was used as a control. Since SynPBS are hemi-discoidal with the tricylindrical cores while GviPBS has a pentacylindrical core, we also added OCP representatives of all clades from cyanobacteria with pentacylindrical PBS cores (as assumed from the alignment of their core-type defining ApcE linkers) as controls (Fig. S18). These controls included *Anabaena* sp. ATCC 29413 (AnaOCP1), *Crinalium epipsammum* PCC 9333 (CrinOCP2), *Gloeocapsa* sp. PCC 7428 (GcapOCP2) and *G. kilaueensis* (GkilOCP3a). These OCPs are all known to quench SynPBS upon photoactivation (*35*, *37*, *38*). However, under similar conditions, we found that SynOCP1, AnaOCP1, CrinOCP2 and GcapOCP2 did not affect the fluorescence intensity, maximum position, or fluorescence lifetime of GviPBS (Fig. 4E, Fig. S19). This indicates that they lost the ability to quench the bundle-shaped PBS during evolution.

Upon photoactivation of GviOCP3a_α, robust quenching of GviPBS fluorescence was observed, which substantially reduced fluorescence of the AP core (∼1.9 ns), but not the components responsible for EET from PE to PC (∼70 ps) and from PC to AP (∼165 ps) (Fig. 4C,D and Fig. S17). Consequently, the GviPBS fluorescence maximum shifted from 663 nm (AP) to 646 nm (PC), indicating GviOCP3a_α binding to the AP core (Fig. 4D,G, Fig. S19). GkilOCP3a, which belongs to a different *Gloeobacter* species, induced similar efficient quenching of GviPBS (Fig. 4E, Fig. S19). Interestingly, both GviOCP3a_α and GkilOCP3a strongly quenched SynPBS as well, just like SynOCP1 did (Fig. S16), suggesting that they share a conserved binding site within the tricylindrical SynPBS core, which is located on the top B cylinder (*7*) (Fig. S14). This Site 1 must be partially preserved in GviPBS to explain its efficient quenching by GviOCP3a_α and GkilOCP3a.

In contrast, GviPBS quenching by GviOCP3a_β was slower and less efficient than by GviOCP3a_α (Fig. 4E,F). This resulted in a partially blue-shifted GviPBS fluorescence maximum (Fig. S19) and unusually altered fluorescence lifetimes (Fig. S16,S17). The fluorescence lifetime of the quenched PBS state may reflect the geometry of the PBS-OCP complex and the mutual orientation of the bilin and the carotenoid (*35*, *37*, *39*), which likely contributes to the observed differences in GviPBS quenching by GviOCP3a_α and GviOCP3a_β. Notably, neither protein affected fluorescence of individual PC and AP samples upon photoactivation (Fig. S19), which underscores the importance of PBS integrity in OCP binding. Nevertheless, our data indicate that GviOCP3a_β accesses different populations of AP chromophores in the core than GviOCP3a_α does, suggesting the existence of a binding site distinct from Site 1. While searching for it, we realized that the OCP footprint at Site 1 on the top B cylinder is also preserved in C/C’ cylinder at the junction of two α and one β AP chains (Fig. S14). This tentative Site 2 is spatially separated from Site 1, allowing noncompetitive OCP binding.

To distinguish this from competition between GviOCP3a_α and GviOCP3a_β for overlapping sites, we exploited the markedly different quenching efficiencies of these two proteins. We titrated GviPBS with each protein, empirically determining the OCP/PBS ratio that led to saturated binding. Then, we compared GviPBS quenching at twice the concentration of either OCP, as well as at an equivalent combined concentration of both. This experiment clearly revealed an additive effect of GviOCP3a_α and GviOCP3a_β, which supports our hypothesis and suggests that the altered efficiency of GviOCP3a_β is due to its binding at Site 2 (Fig. 4F,G and Fig. S14). The location of Site 2 on C/C’ cylinders, which are unique to PBS cores containing “3+2” cylinders, emphasizes the importance of C/C’ cylinders as independent EET pathways and as targets for the regulation by GviOCP3a_β. Interestingly, while GviOCP3a_α and GkilOCP3a induced strong quenching of SynPBS lacking C/C’ core cylinders, GviOCP3a_β was completely dysfunctional on SynPBS (Fig. S16). This finding corroborates the existence of multiple binding sites in GviPBS and potentially in other PBS with pentacylindrical cores. We hypothesize that GviOCP3a_α is the most ancestral extant OCP, because it is highly effective on both GviPBS and SynPBS, acting at Site 1. We also hypothesize that GviOCP3a_β is a relatively new type of OCP whose duplication and subsequent evolution enabled the regulation of EET pathways over C/C’ cylinders by binding at Site 2.

In summary, we conclude that the additional C/C’ cylinders are a unique structural feature of the pentacylindrical core that adds extra EET pathways to A/A’ cylinders. In turn, this requires additional solutions to regulate these energy flows. In this regard, having two sites for different OCPs seems to be a reasonable evolutionary solution, which may also be important for preventing energy migration between neighboring cores, while allowing for targeted energy transfer from rods to A/A’ cylinders to fuel the photosynthesis.

## Acknowledgements

The authors are thankful to Prof. Alexey Amunts for help with preliminary screening of GviPBS samples on grids, to Dr. Rustam H. Ziganshin for help with running LC-MS for identification of GviPBS proteins and to Daniil M. Panin for help with OCP data analysis.

We thank Resource Center of Probe and Electron Microscopy of National Research Center ‘‘Kurchatov Institute’’ (http://rc.nrcki.ru/pages/main/nanozond/facilities/12604/index.shtml). This work has been carried out using computing resources of the federal collective usage center Complex for Simulation and Data Processing for Mega-science Facilities at National Research Center ‘‘Kurchatov Institute’’ (http://ckp.nrcki.ru).

## Author contributions

Conceptualization: NNS

Methodology: ADB, YBS, TNB, EGM, VOP, NNS

Investigation: ADB, YBS, TNB, MAS, DAG, GVT, DAC, EGM, KMB, NNS

Visualization: ADB, YBS, EGM, NNS

Funding acquisition: MAS, VOP, KMB, NNS

Project administration: NNS

Supervision: KMB, NNS

Writing – original draft: NNS

Writing – review & editing: NNS, with input from all authors

## Data availability

The atomic coordinates have been deposited in the Protein Data Bank with the accession codes 9V7J (PBS core), 9V7K (PBS C), 9V7G (PBS Rx), 9V7L (PBS R1), 9V7H (PBS R2) and 9V7I (PBS R3). The cryo-EM maps have been deposited in the Electron Microscopy Data Bank with accession codes EMD-64815 (PBS core), EMD-64816 (PBS C), EMD-64812 (PBS Rx), EMD-64817 (PBS R1), EMD-64813 (PBS R2), EMD-64814 (PBS R3), EMD-64828 (consensus), EMD-64868 (composite), EMD-64851 (diverging rods) and EMD-64852 (parallel rods). Raw time-resolved emission spectra for GviPBS in the absence or in the presence of GviOCP3a_α, GviOCP3a_β, or their combination have been uploaded to Zenodo (https://doi.org/10.5281/zenodo.15575709). Other data are available in the paper or in Supplementary information.

## Supplementary materials

### Materials and Methods

#### Phycobilisome isolation

The strain of *Gloeobacter violaceus* PCC 7421 used in this study was provided by the Collection of microalgae and cyanobacteria IPPAS of the K.A. Timiryazev Institute of Plant Physiology of the Russian Academy of Sciences (collection ID: IPPAS-B470). *G. violaceus* culture was grown without shaking for 2 months in 1 L Erlenmeyer flasks containing 0.5 L of standard BG-11 medium (*40*) buffered with 10 mM HEPES at 22 °C and illuminated with LED (4000 K, 8W) with light intensity 10 μmol photons×s^−1^×m^−2^. The cells were harvested by centrifugation at 3300×g for 30 min and washed once in 0.65 M potassium phosphate buffer (pH 7.4) with 0.5 M sucrose, centrifuged and suspended in 6 ml of the same buffer to a concentration of 0.1 g (wet weight) per ml. The cells were disrupted through a french press at 1500 psi (Thermo electron) three times. Triton X-100 was added to the brown lysate to the final concentration of 2% (v/v), and the solution was gently shaken for 20 min under aluminium foil. Unbroken cells and cell debris were removed by centrifugation at 15557×g for 45 min. The blue-violet PBS containing supernatant fraction (830 µl per tube) was loaded onto a step sucrose gradient: 0.6 mL of 2.0 M; 2 mL of 1.0 M; 2 mL of 0.75 M, 2 mL of 0.6 M, and 2.4 ml of 0.5 M sucrose made on 0.75 M potassium phosphate buffer (pH 7.4). The resulting gradients were centrifuged at 288000×g for 6 h using a SW41 Ti rotor on a Beckman L8-80M centrifuge at 25 °C. All buffers contained 3 mM sodium azide and procedures described above were carried out at 25 °C. Intact phycobilisomes (PBS) were recovered from the middle part of the 1 M sucrose layer. Collected samples were stored at 4 °C for immediate use. Long-term preservation (at least 3-4 months) could be achieved by adding ammonium sulfate until the final concentration of 1 M.

#### Phycobiliprotein separation

Cells were harvested as described above and frozen at –80 °C until needed. Cell pellet was thawed and washed twice with a buffer A (30 mM Tris-HCl (pH 8.1), 5 mM EDTA) and suspended in the same buffer containing 0.1 mM PMSF and 0.26 mg/ml lysozyme, to a concentration of 0.4 g (wet weight) per ml. The cells were disrupted through a french press at 1500 psi (Thermo electron, USA) three times. Brown lysate was centrifuged at 52360×g for 30 min using LG-25M centrifuge (Shuke Instrument, China) at 4 °C to remove cell debris and membrane fragments. The resulting supernatant was fractionated with ammonium sulfate. Solid ammonium sulfate was added to 20% of saturation and the turbid solution was centrifuged at 52360×g for 30 min, which led to partial phycobiliprotein precipitation. Ammonium sulfate was added to 40% saturation and centrifugation was repeated, yielding deep blue supernatant and precipitation of pink protein, which was resuspended in buffer A. Ammonium sulfate was raised to 60% of saturation. Centrifugation was repeated and the blue precipitate was resuspended in buffer A. Pink and blue fractions from 40% and 60% ammonium sulfate saturation, respectively, were dialyzed against 10 mM Tris-HCl pH 7.85 at 4 °C overnight. The dialyzed solutions of pink and blue fractions were clarified by centrifugation at 52360×g for 30 min. Both fractions were further purified in the same way by anion-exchange chromatography on a HiTrapQ HP column (5 ml, Cytiva) as follows. The protein sample was loaded on the column pre-equilibrated with the dialysis buffer, and washed with 20 mM Tris-HCl (pH 7.6) until the flowthrough fraction became completely colorless. Then, bound phycobiliproteins were eluted with a linear gradient of 0-0.5 M NaCl in 20 mM Tris-HCl (pH 7.6). Phycoerythrins from the pink fraction eluted at 140-250 mM NaCl, whereas phycocyanins and allophycocyanins from the blue fraction eluted at 110-260 mM NaCl and 320-500 mM NaCl, respectively. Phycobiliprotein preparations were stored at 4 °C.

#### GraFix

PBS have a high tendency to dissociate in low-ionic strength buffers and are often chemically stabilized for structural studies (*5*, *15–17*). We used the GraFix method (*15*) for mild chemical crosslinking of PBS without undesired particle aggregation. The ultracentrifuged PBS fraction was dialyzed against 0.75 M potassium phosphate buffer (pH 7.4) with 0.4 M sucrose at 4 °C overnight and passed through a filter paper. The resulting solution was concentrated with 30-kDa Vivaspin Turbo 4 centricon (Sartorius, Germany) till the final sample absorbance at 566 nm equal to 4.9. The concentrated sample (150 µl per tube) was loaded onto a step sucrose (M); glutaraldehyde (GA) (v/v %) gradient made with 0.75 M potassium phosphate buffer pH 7.4: 1 mL of 2.0 M; 0.05%; 2 mL of 1.0 M; 0.03%; 2 mL of 0.75 M; 0.01%, 2 mL of 0.6 M; 0.005%, and 2.9 ml of 0.5 M. Control gradient with the PBS sample was made without GA to exclude its adverse effect on the sedimentation pattern. Centrifugation was performed at 288000×g for 6 h using a SW41 Ti rotor on a Beckman L8-80M centrifuge at 25 °C. After centrifugation, crosslinked PBS (GraFix-PBS) fractions were obtained from the 1 M sucrose layer completely separated from the aggregated PBS. Isolated GraFix-PBS fraction was quenched with glycine-KOH pH 7.7 (final concentration 73 mM) and dialyzed against 50 mM Tris-HCl (pH 7) at 4 °C overnight. GraFix-PBS were concentrated with 30-kDa Vivaspin Turbo 4 centricon (Sartorius, Germany) till final sample absorbance at 566 nm equal to 8.074 and used for grid preparation.

#### Cryo-EM sample preparation

A 3 μl of the sample was applied to the Quantifoil R1.2/1.3 300 mesh Cu grid coated with 2 nm amorphous carbon film, which was not glow-discharged prior to sample application. The grid was plunge-frozen in liquid ethane using a Vitrobot Mark IV (Thermo Fisher Scientific, USA) with the following settings: chamber humidity 100 %; chamber temperature 4 °C; blotting time 3 s; blotting force 0. The grid was then stored in liquid nitrogen until use.

#### Cryo-EM data collection

Cryo-EM data were collected on a Titan Krios 60-300 transmission electron microscope (Thermo Fisher Scientific, USA) equipped with a field emission electron gun X-FEG (Thermo Fisher Scientific, USA), spherical-aberration corrector (CEOS GmbH, Germany), a post-column BioQuantum energy filter (Gatan, USA) and a K3 direct electron detector (Gatan, USA) in counting non-CDS mode using SerialEM 4.055 and 9-hole image-shift data acquisition strategy at the National Research Centre ‘‘Kurchatov Institute’’. The microscope was operated at 300 kV with a nominal magnification of 81,000x, corresponding to a pixel size of 0.874 Å at the specimen level, and an electron energy selecting slit of 20 eV. A total dose of 66 e^−^/Å^2^ within a 3.9 s exposure time was fractionated into 60 frames, resulting in an electron dose of 1.1 e^−^/Å^2^ per frame. In total, ∼45,000 movies were collected in a nominal defocus range from −0.7 to −1.7 mm with a step of 0.1 mm. Detailed parameters of data acquisition are listed in Table S2.

#### Cryo-EM data processing

The initial cryo-EM data were pre-processed in Warp ver. 1.0.9 (*41*). Global and local motion estimation, CTF model estimation, particle picking using a specifically re-trained deep CNN BoxNet were performed for all the movies. All images were inspected in a semiautomated manner using Warp’s thresholds for defocus (less than 2 mm), estimated resolution (less than 4 Å) and drift (less than 2 Å). A total of 1,364,249 particles were picked from 41,564 selected images and extracted into 960 pixel boxes. The motion-corrected particle stacks were imported into CryoSPARC ver. 4.7 (*42*) for further data processing. Several rounds of 2D classification were performed using downsampled to 480 pixels boxes, 50 classes at each step to maximize the number of true-positive particles in the subset of interest. Reference-free 2D classification revealed an accurate C2 symmetry of the PBS particles. A cleaned subset of 746,972 PBS particles, downsampled to 672 pixels (1.25 Å per pixel), was used for *ab initio* initial map reconstruction and homogeneous refinement. Following rounds of refinement with a C1 and C2 symmetry, per-particle defocus adjustment, and adjustments to the higher-order CTF terms for each exposure group, as well as local refinement of the most stable core part of the PBS, a 2.85 Å resolution cryo-EM density map of the PBS core was obtained (EMD-64815).

To elucidate the structural intricacies of the more dynamic parts of the PBS as well as the structure of the AP core hexamers, five soft masks were created using UCSF Chimera (*42*) based on the locally refined PBS core map. Further local refinement resulted in two maps of the core part: 2.94 Å PBS C (EMD-64816), 2.72 Å PBS Rx (EMD-64812), and three maps of the hexamers of the 1st and the 2nd rods: 3.76 Å PBS R1 (EMD-64817), 2.72 Å PBS R2 (EMD-64813), 3.03 Å PBS R3 (EMD-64814).

To elucidate the structural heterogeneity of the rods, signal subtraction of the core part was performed followed by heterogeneous refinement. Two distinct conformations with diverging and parallel rods were obtained. Subsequent non-uniform refinement yielded a 4.25 Å map for diverging conformation (EMD-64851) and a 4.66 Å map for parallel conformation, with C2 and C1 symmetry imposed, respectively. Schematic representation of data processing is shown in Fig. S2.

#### Model building and refinement

Crystal structures of AP, PC and PE hexamers from *G. violaceus* (PDB codes: 2vjt, 2vjr and 2vjh, respectively) as well as cryo-EM structures of the ApcE and ApcC proteins from *Anabaena* 7120 (PDB 7eyd) were used for the initial modeling. The atomic models were first docked into the corresponding cryo-EM density maps using UCSF ChimeraX (*43*). The conformational differences in hexamers were manually adjusted in COOT (*44*). Sequences of the linker proteins were fetched from Uniprot – ApcC (Q7NL78), ApcE (Q7NL81), ApcF (Q7NJA2), Glr1262 (Q7NL64), Glr2806 (Q7NGT2), CpcC1 (Q7NM19), CpcC2 (Q7NGF2).

Two ApcF and eight ApcC were built using “mutate residue range” to the corresponding subunits. ApcE was manually adjusted and mutated from the template model. Linker proteins Glr1262, Glr2806, CpcC1 and CpcC2 were built *de novo* by tracing the backbone, assigning the sequence and fitting the side chains. CpcC1 and CpcC2 sequences were extended in N-termini according to the previous work (*22*). All the structures were modelled and fitted using COOT (*44*). The final atomic models were refined in real space using Servalcat Refmac in CCPEM and geometric restraints (*45*). The final atomic models were evaluated using Molprobity (*44*) and the statistics of data collection and model validation were included in Table S2.

#### Cloning, expression and purification of OCP

cDNA corresponding to OCP3a homologs of *G. violaceus* PCC 7421 was obtained by PCR amplification from its genome using gene-specific primers GviOCP3a_α_forw (5’-ATATACATATGGCGTTTACCCTTGAATC-3’), GviOCP3a_α_rev (5’-TTATTCTCGAGTTACCGGTAGCTGGGAG-3’), GviOCP3a_β_forw (5’-ATATACATATGGTACTCACCATCGAGTC-3’), GviOCP3a_β_rev (5’-TTATTCTCGAGCTAACGGGCTAGCTTGTCC-3’). Primers contained *Nde*I and *Xho*I restriction sites for subsequent cloning into the pET28_his_3C vector (kanamycin resistance). To obtain holoforms of OCP3a proteins with the embedded echinenone (ECH), the OCP plasmids were transformed into C41(DE3) *E. coli* cells carrying the pACCAR25ΔcrtXZcrtO plasmid (chloramphenicol resistance) which enabled ECH synthesis (*46*). Protein expression was induced by adding 0.1 mM isopropylthiogalactoside and continued for 24 h at 27 °C. Protein purification included immobilized metal-affinity chromatography (IMAC), subtractive IMAC after His-tag cleavage by rhinovirus 3C protease and gel-filtration on a Superdex 200 26/600 column (GE Healthcare). After His-tag removal, extra N-terminal GPH residues are left due to plasmid design. Electrophoretically homogeneous preparations were obtained. Typical Vis/UV absorbance ratios for OCP3 samples obtained were in the 1.7-1.8 range, indicating the lack of apoprotein (*35*, *37*). Purified proteins were stored at −80 °C. The oligomerization of proteins was characterized by analytical size-exclusion chromatography (SEC) on a Superdex 200 Increase 5/150 column (Cytiva) as described earlier (*35*, *37*). The absorbance spectra of GviOCP3a_α and GviOCP3a_β were recorded online during the SEC runs using Agilent 1260 II Infinity chromatography system equipped with the diode-array detector.

SynOCP1 and AnaOCP1 (*38*), CrinOCP2 and GcapOCP2 (*37*), as well as GkilOCP3a (*35*) proteins were obtained exactly as described in previous works.

#### HPLC

Carotenoid content in *G. violaceus* cells and the recombinant proteins was analyzed by high-performance liquid chromatography (HPLC) on a Nucleosil C18 column 4.6*250 mm at a flow rate of 1 ml/min and temperature set at 28 °C as described earlier (*37*). Carotenoid extraction was performed by acetone with subsequent partitioning with n-hexane. 100 μL of purified protein or 20 mg of cell precipitate were mixed with 100 μL acetone and vortexed thoroughly until full destruction of cells or protein denaturation. Then 100 μL of hexane was added and the resulting mixture was vortexed and centrifuged for phase separation. The upper hexane fraction was dried under nitrogen, redissolved in 25 μL acetone and loaded onto a column. Peak positions and spectra were compared with standards (canthaxanthin, echinenone and β-carotene) for identification of carotenoids in the sample.

#### Photoactivity assay

To measure the photocycle of GviOCP3a homologs (5 μM) in 20 mM Tris-HCl buffer, pH 7.6, containing 150 mM NaCl and 0.1 mM EDTA, the samples were transferred to a 1-cm quartz cuvette and the absorbance spectrum was continuously recorded. A blue light-emitting diode (M455L3, Thorlabs, USA) with a maximum emission at 445 nm was used to photoswitch OCP to its light-adapted state. Temperature was maintained by a Peltier controlled cuvette holder Qpod 2e (Quantum Northwest, USA). After reaching the equilibrium, the blue LED was switched off and subsequent transitions to the dark-adapted state were followed by changes of absorbance at 550 nm. For quantitative comparison of GviOCP3a homologs, we built an Arrhenius plot (*35*, *37*). Kinetics of the R-O transition in the temperature range 5–25 °C were approximated by exponential decay and the corresponding time constant (t) for each temperature was derived. ln(1/t) (rate constant) against 1000/T (K) was linear and was used to derive activation energy (E_a_).

Absorption spectra of *G. violaceus* cells were recorded using a stabilized white light source with an SLS204 deuterium lamp (Thorlabs, USA) and CCD spectrometer Maya2000Pro (Ocean Optics, USA) coupled with an integrating sphere (Tarusov Electrics, Russia).

#### PBS quenching

Steady-state fluorescence spectra of isolated PBS and PBPs were measured with a plate reader CLARIOstar^plus^ (BMG Labtech, Germany). All measurements were performed in a black 384 well microplate with flat bottom (Greiner Bio-One, Germany) using 0.75 M potassium phosphate (pH 7.4) or 20 mM Tris-HCl (pH 7.6) buffers for PBS and PBPs, respectively. Typically, fluorescence was excited at 575 nm with an emission range of 600-800 nm. In case of PE preparation, the excitation wavelength was changed to 475 nm for determination of its full emission spectra at 500-800 nm. Fluorescence quenching with different OCP homologs in microplate wells were monitored before and after photoactivation with a blue light-emitting diode (M455L3, Thorlabs, USA) with a maximum emission at 445 nm for 2 min at 25 °C.

To measure time-resolved PBS fluorescence, a setup based on the TOPOL-1050-C optical parametric oscillator (Avesta Project LTD, Moscow, Russia) was used. The sample was excited by a sequence of 150-fs laser pulses at 80 MHz repetition rate. The infrared laser emission (970 nm) was converted to the visible range (485 nm) using a second-harmonic generator. The average laser power was set to 3 mW. For wavelength selection of the detected signal, an ML-44 monochromator (Solar LS, Belarus) was used. A longpass filter (FEL0600, Thorlabs, USA) was placed in front of the detector. Picosecond fluorescence decay kinetics and changes in total signal intensity were recorded in time-correlated single-photon counting (TCSPC) mode using a cooled HPM-100–07 C detector (Becker & Hickl, Germany). To record fluorescence intensity kinetics in the 1–1000 s range, the FIFO mode of the SPCM software package (Becker & Hickl, Germany) was used. For time-resolved emission spectroscopy (TRES) in the 500–750 nm range, a combination of the Gemini interferometer (Nireos, Italy) and a cooled HPM-100–40 C detector (Becker & Hickl, Germany) was used. For photoactivation of OCP and recording NPQ kinetics of PBS, a blue LED (SOLIS-445C, Thorlabs, USA) was used, with its emission further filtered by a bandpass filter (450 nm, 40 nm width, Thorlabs, USA). The LED power was maintained at 500 mW in all experiments. The temperature of the 500 µL PBS solution in an acrylic cuvette was stabilized at 10 °C using a Qpod 2e temperature controller (Quantum Northwest, USA) to prevent spontaneous fluorescence recovery.

Global analysis fitting was performed for the time-resolved fluorescence spectra using the Matlab (USA) software. With global analysis, all wavelengths were analyzed simultaneously with a set of common time constants. An experimentally measured instrumental response function (IRF) was used to approximate the fluorescence decay kinetics and to determine the characteristic lifetimes. For this purpose, we recorded the signal from the laser as Raman scattering of water. The kinetic parameters were found by numeric nonlinear minimization of the solution to fit the experimental data in the spectral interval of 500–750 nm and the time range of 10 ps–12.5 ns. The post-processing and visualization of calculated data were performed using the OriginPro 2024 package (OriginLab Corp., USA) essentially as described earlier (*47*).

Effect of combination of different OCPs on GviPBS quenching was analyzed using Jin’s formula: Q = E(α+β) / (Eα + Eβ - Eα x Eβ) (*48*). Here, Eα, Eβ and E(α+β) are average effects (relative reduction of the nanosecond component) of individual OCP photoactivation, or their combination, respectively. In this method, Q < 0.85 indicates antagonism; 0.85 ≤ Q ≤ 1.15 indicates additive effects; Q ≥ 1.15 indicates synergism.

#### Phylogenetic analysis

Construction of the phylogenetic tree was carried out for OCP, Glr2806 and Glr1262 homologs using the same protocol, which is detailed using the OCP case as follows. Protein sequences of full-length OCP homologs (OCP1, OCP2 and OCP3 representatives) were retrieved from the NCBI core nucleotide BLAST database (GenBank+EMBL+DDBJ+PDB+RefSeq), update date:2025/05/25. Multiple sequence alignment was performed using Clustal Omega in the EMBL-EBI Job Dispatcher sequence analysis tools framework (*49*). A tree was built using MEGA (*50*) using Maximum Likelihood method and Jones-Taylor-Thornton model (*51*) of amino acid substitutions. The analytical procedure encompassed 146 sequences. Gap positions in the alignment were completely deleted resulting in a final data set comprising 302 positions. The initial tree for the heuristic search was selected by choosing the tree with the superior log-likelihood between a Neighbor-Joining (NJ) tree (*52*) and a Maximum Parsimony (MP) tree. The NJ tree was generated using a matrix of pairwise distances computed using the Jones-Taylor-Thornton model (*51*). The MP tree had the shortest length among 10 MP tree searches, each performed with a randomly generated starting tree. The number of bootstrap replicates was determined adaptively (*50*). Family clades were supported by a high percentage (>90 % of 118 replicates) of trees in which the associated taxa clustered together. According to our criterion, 20 representatives of OCP3c are not clustered reliably either with OCP3 or other OCP groups and included into the OCP3 group based on previous studies (*30*, *35*).

### Supplementary Tables

**Table S1.** Mass spectrometry-based identification of PBS proteins from *G. violaceus*.

| Protein | Category | NCBI Reference | Uniprot ID | Mw, Da | Sequence coverage (MALDI) | Expect value (MALDI) | Sequence coverage (LC-MS) |
| --- | --- | --- | --- | --- | --- | --- | --- |
| ApcA | PBPs | WP_011141246.1 | Q7NL80 | 17537.1 | 59% | 2.6e-004 | 86.3% |
| ApcB | PBPs | WP_011141247.1 | Q7NL79 | 17211.7 | - | - | 91.3% |
| ApcC | linkers | WP_011141248.1 | Q7NL78 | 7754.0 | 52% | 1.1e-008 | 52.2% |
| ApcD | PBPs | WP_011141181.1 | Q7NLE3 | 18032.8 | - | - | 87.6% |
| ApcE | PBPs/linkers | WP_011141245.1 | Q7NL81 | 129836.6 | 57% | 2.8e-055 | 72.1% |
| ApcF | PBPs | WP_011141928.1 | Q7NJA2 | 17433.9 | - | - | 80.1% |
| CpcA | PBPs | WP_011141185.1 | Q7M7F7 | 17662.9 | 82% | 1.4e-005 | 94.4% |
| CpcB | PBPs | WP_011141184.1 | Q7M7C7 | 18459.9 | 79% | 2.8e-012 | 97.7% |
| CpcC1 | linkers | WP_164928640.1 | Q7NM19 | 31043.1 | 77% | 3.5e-014 | 77.2% |
| CpcC2 | linkers | WP_231848246.1 | Q7NGF2 | 30877.9 | 65% | 6.0e-006 | 66.2% |
| CpcD-1 | linkers | WP_011141267.1 | Q7NL59 | 7769.9 | 24% | 1.4e-011 | 90.0% |
| CpcD-2 | linkers | WP_011141266.1 | Q7NL60 | 8169.4 | 48% | 4.5e-007 | 72.4% |
| CpeA | PBPs | WP_011141189.1 | Q7NLD7 | 17658.0 | 59% | 3.1e-004 | 94.5% |
| CpeB | PBPs | WP_011141190.1 | Q7NLD6 | 18427.2 | 33% | 2.0e-002 | 83.6% |
| CpeC | linkers | WP_011141263.1 | Q7NL63 | 31825.2 | 75% | 2.2e-013 | 87.4% |
| CpeD | linkers | WP_011141264.1 | Q7NL62 | 28400.2 | 81% | 8.9e-014 | 67.8% |
| CpeE | linkers | WP_011141265.1 | Q7NL61 | 28354.9 | 53% | 1.4e-010 | 66.9% |
| Glr1262 | linkers | WP_011141262.1 | Q7NL64 | 92019.8 | 53% | 5.6e-032 | 74.9% |
| Glr2806 | linkers | WP_011142800.1 | Q7NGT2 | 81422.5 | 44% | 1.1e-007 | 43.2% |

Red font indicates polypeptides not covered by MALDI MS, but reliably confirmed by LC-MS.

**Table S2.**
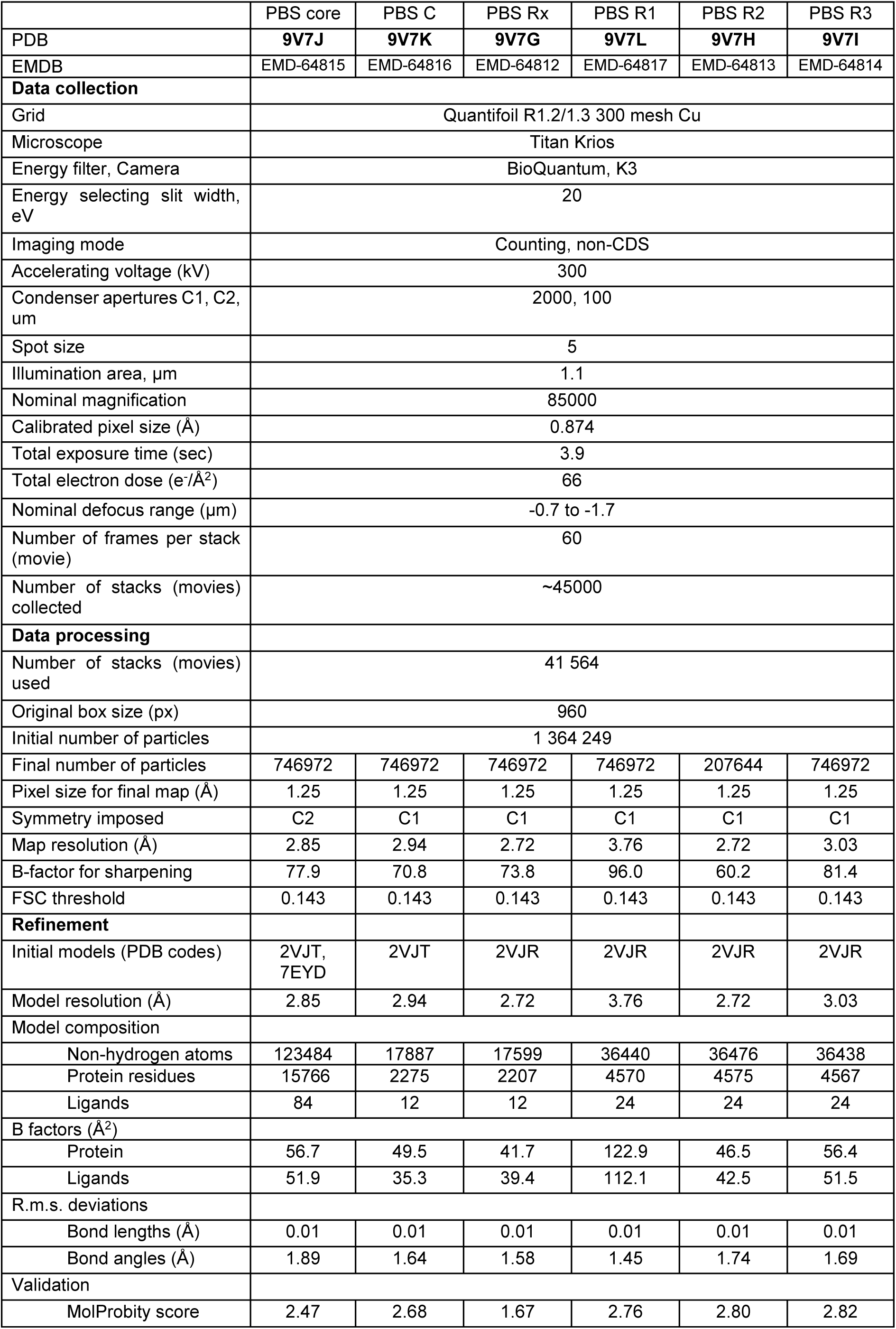
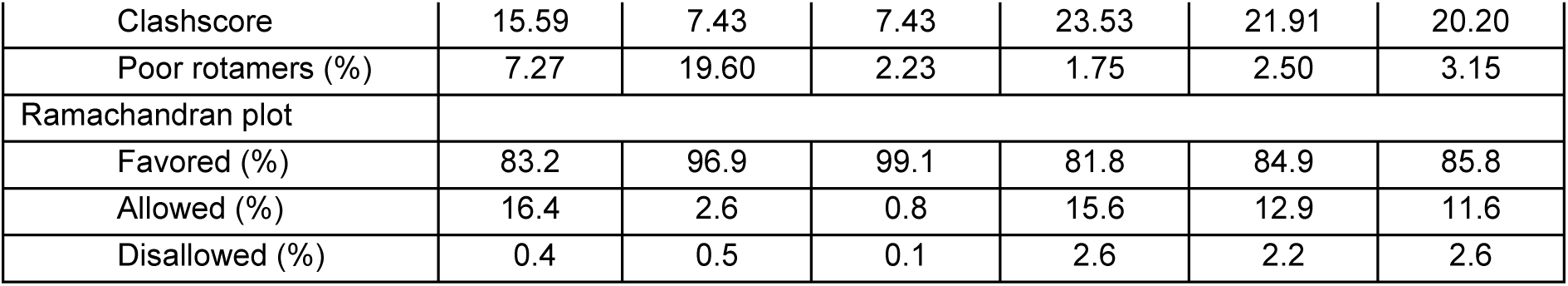
Cryo-EM data collection, refinement and validation statistics.

### Supplementary figures

**Fig. S1.**
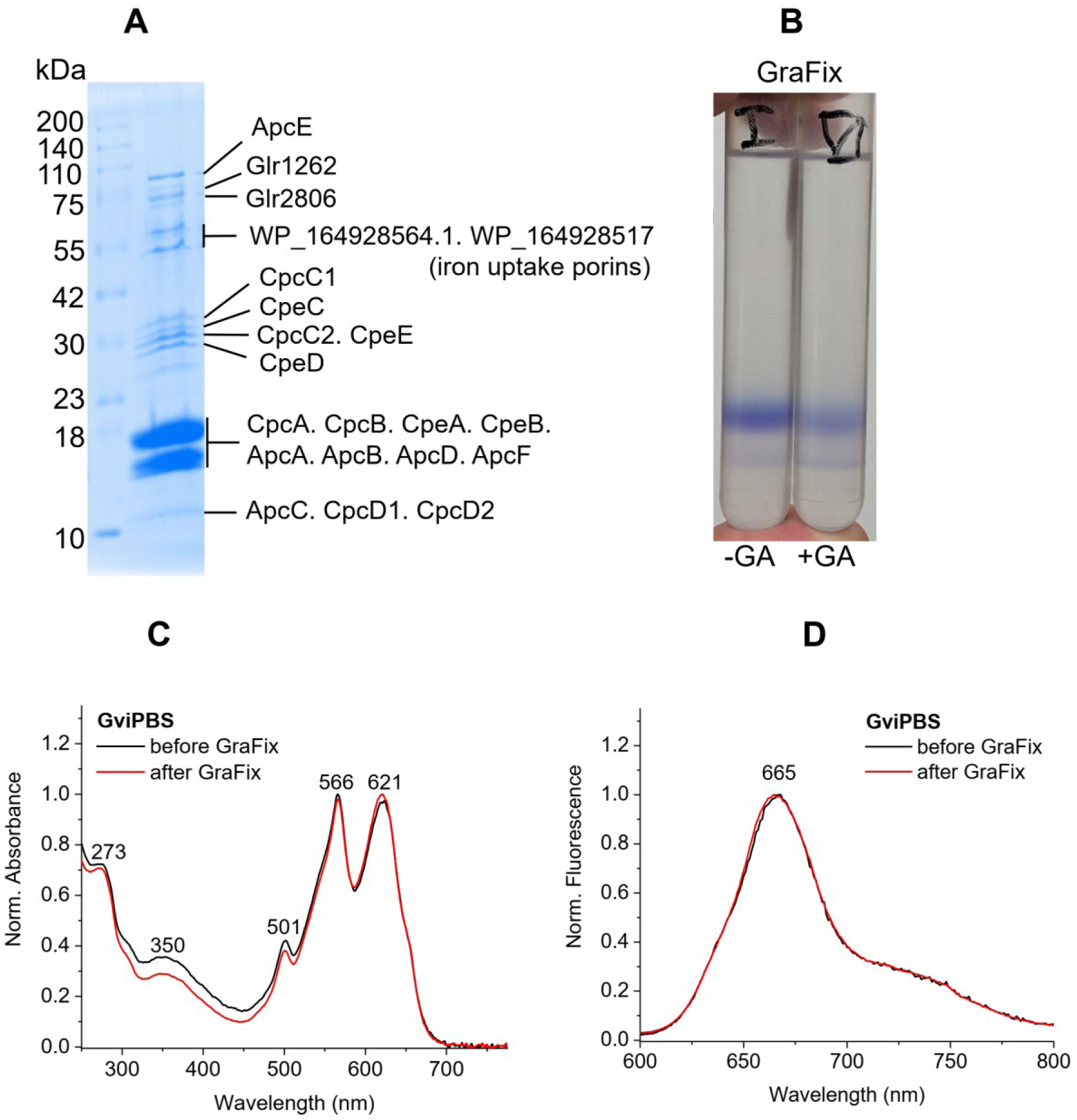
*G. violaceus* PBS purification and characterization. **A**. SDS-PAGE analysis of the protein constituents of GviPBS. **B**. Ultracentrifugation tubes showing that the sedimentation pattern of GviPBS did not change in the presence of glutaraldehyde (GA) during GraFix (*15*). Absorbance (**C**) and fluorescence (**D**) spectra of the GviPBS sample obtained by ultracentrifugation before and after GraFix showing the preservation of the GviPBS assembly.

**Fig. S2.**
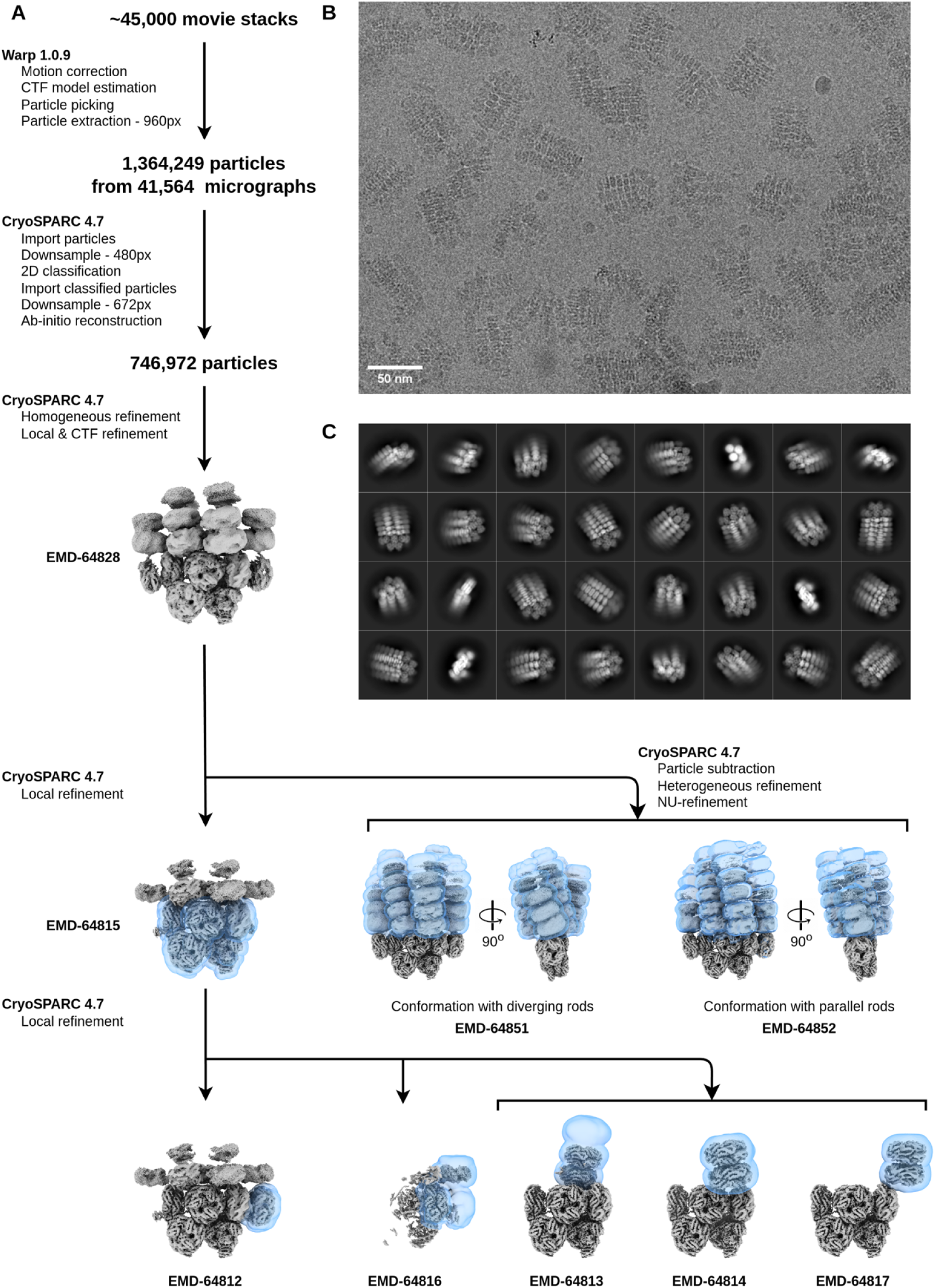
Cryo-EM data processing workflow and 3D reconstruction. **A**. The workflow for the cryo-EM data processing. Cryo-EM density maps of GviPBS are shown in grey, masks used for the local refinement are shown in transparent blue. Each map is labeled with the corresponding EMDB code. **B**. A representative motion-corrected electron micrograph of GviPBS. The scale bar corresponds to 50 nm. **C**. 2D classification of the cryo-EM data demonstrating different views of the GviPBS.

**Fig. S3.**
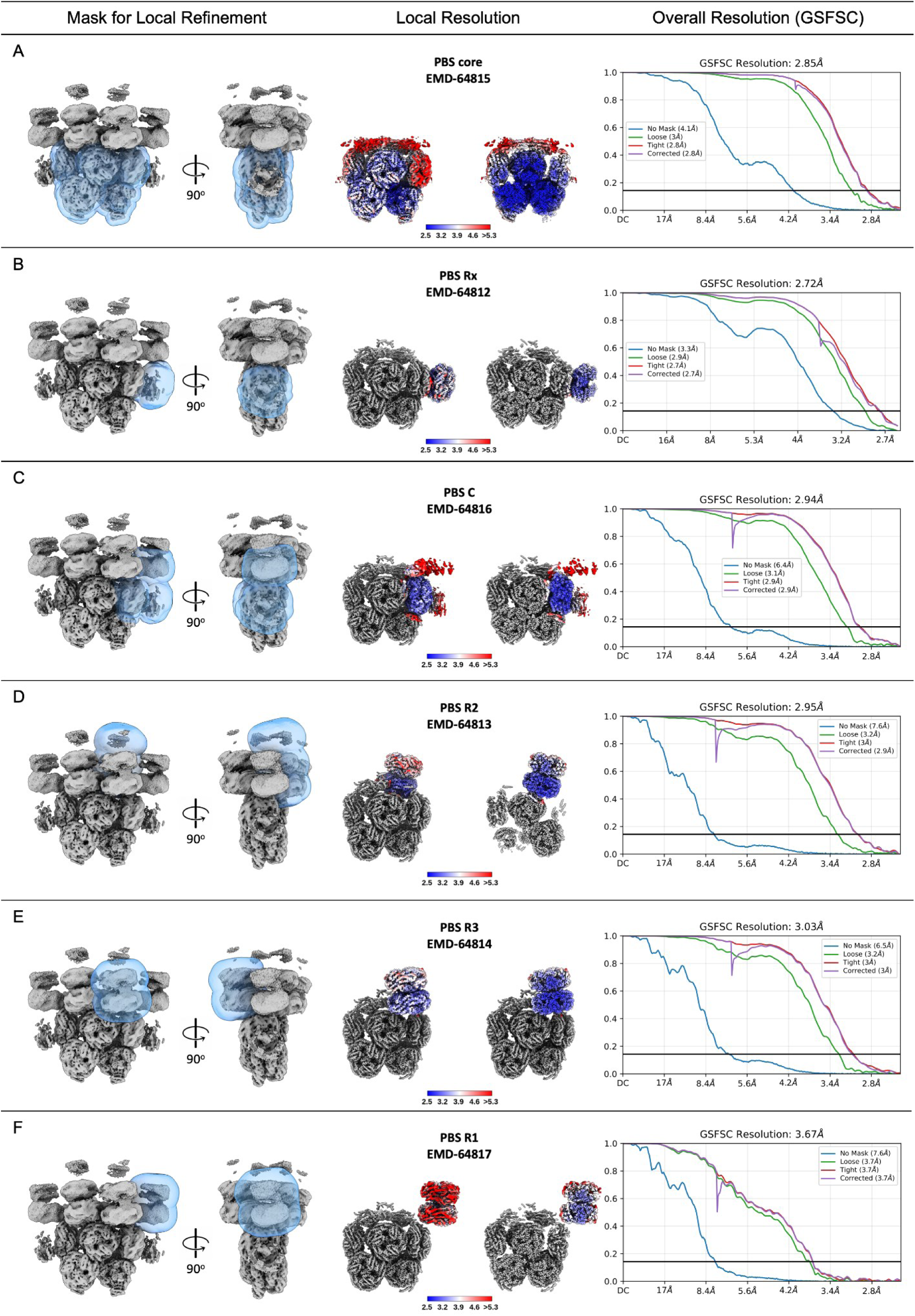
Overall and local resolution estimation for each local refinement. Left panel: The masks used for the local refinement. The consensus cryo-EM density map is shown as gray, and the mask used for local refinement is shown as transparent blue. Middle panel: Local resolution estimation for corresponded parts of PBS. The consensus cryo-EM density map is shown as gray, the locally refined cryo-EM density maps are colored according to the local resolution calculated by cryoSPARC, the color key is in Angstroms. Right panel: Gold-standard FSC plots for the locally refined reconstructions. The global resolutions were estimated by gold-standard FSC (threshold at 0.143).

**Fig. S4.**
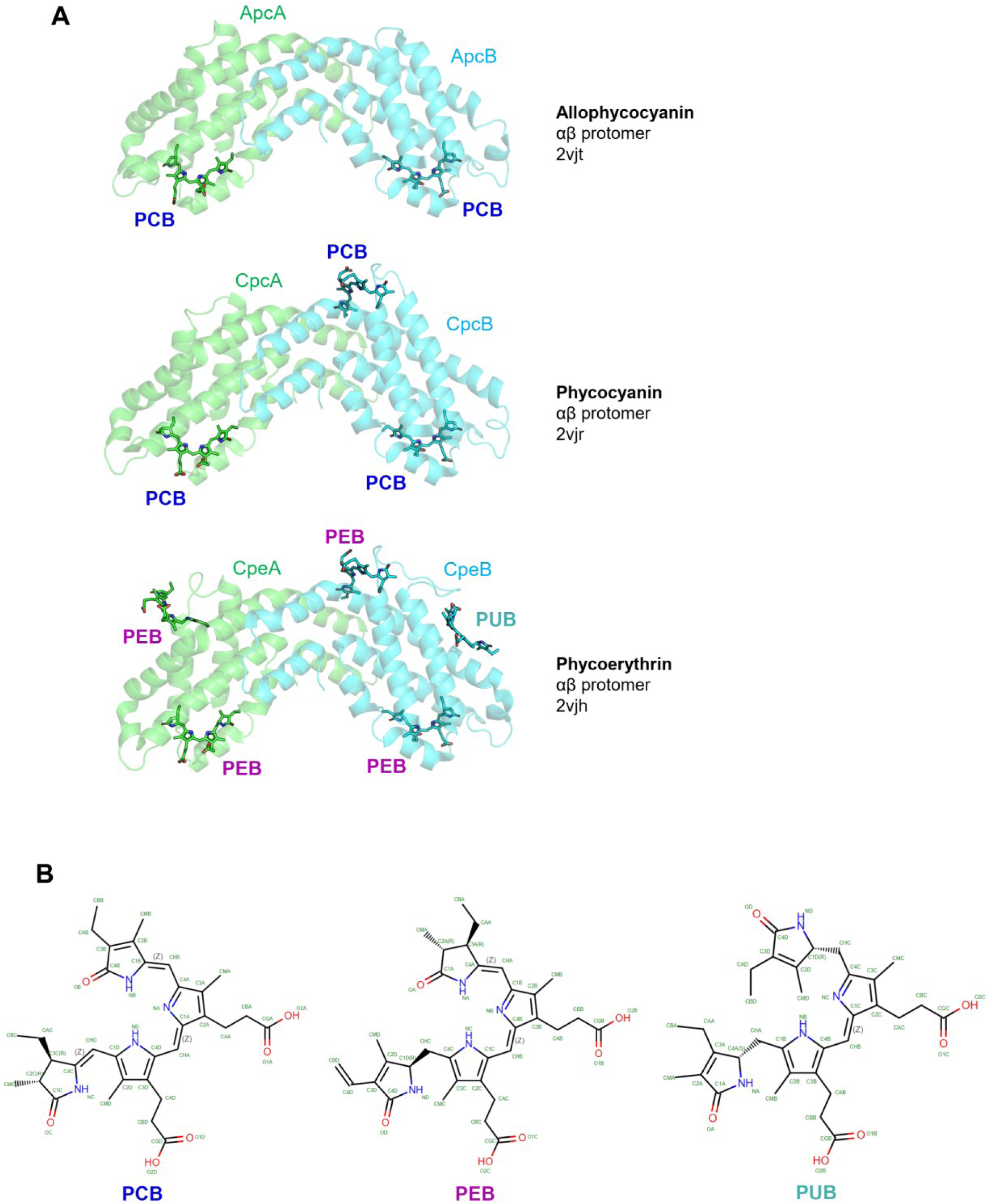
Crystallographic structures of αβ protomers of allophycocyanin (AP), phycocyanin (PC) and phycoerythrin (PE) from *G. violaceus* with three main chromophores covalently attached to these PBPs. PCB – phycocyanobilin, PEB – phycoerythrobilin, PUB – phycourobilin. PDB codes are indicated. **A**. Cartoon representation of the PBP protomers with bilins shown as sticks. **B**. Chemical structures of the bilins from *G. violaceus* PBS.

**Fig. S5.**
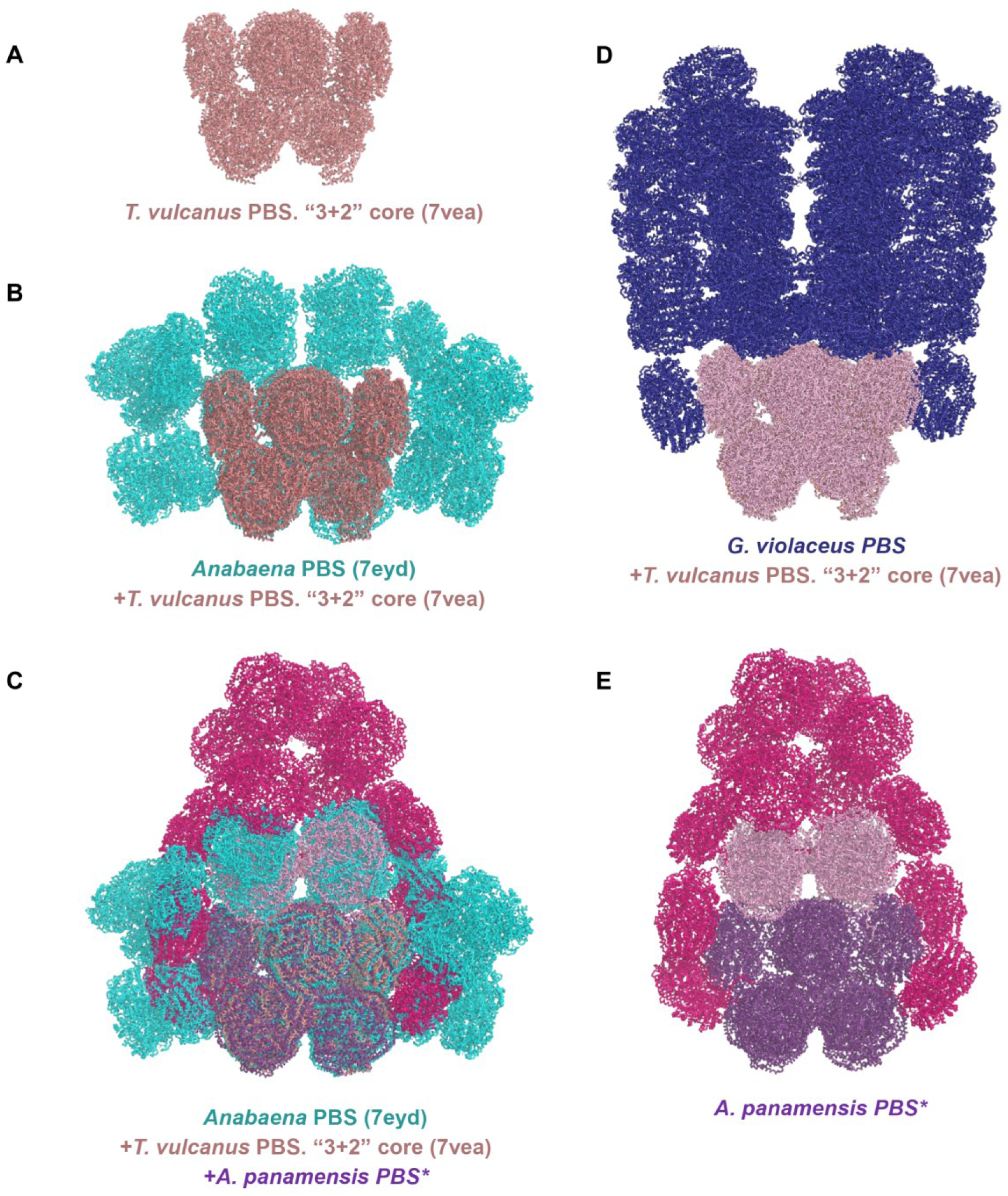
The pentacylindrical core as the common substructure of PBS with different morphologies. **A**. The pentacylindrical core of *T. vulcanus* PBS. **B**. The pentacylindrical core of *Anabaena* PBS (*8*) shown overlaid with that of *T. vulcanus* (*17*). **C**. Overlay of the structures from panel B with the structure of the paddle-shaped PBS from *A. panamensis* (*5*). **D**. GviPBS with the pentacylindrical core (this work). **E**. The paddle-shaped PBS from *A. panamensis* (*5*).

**Fig. S6.**
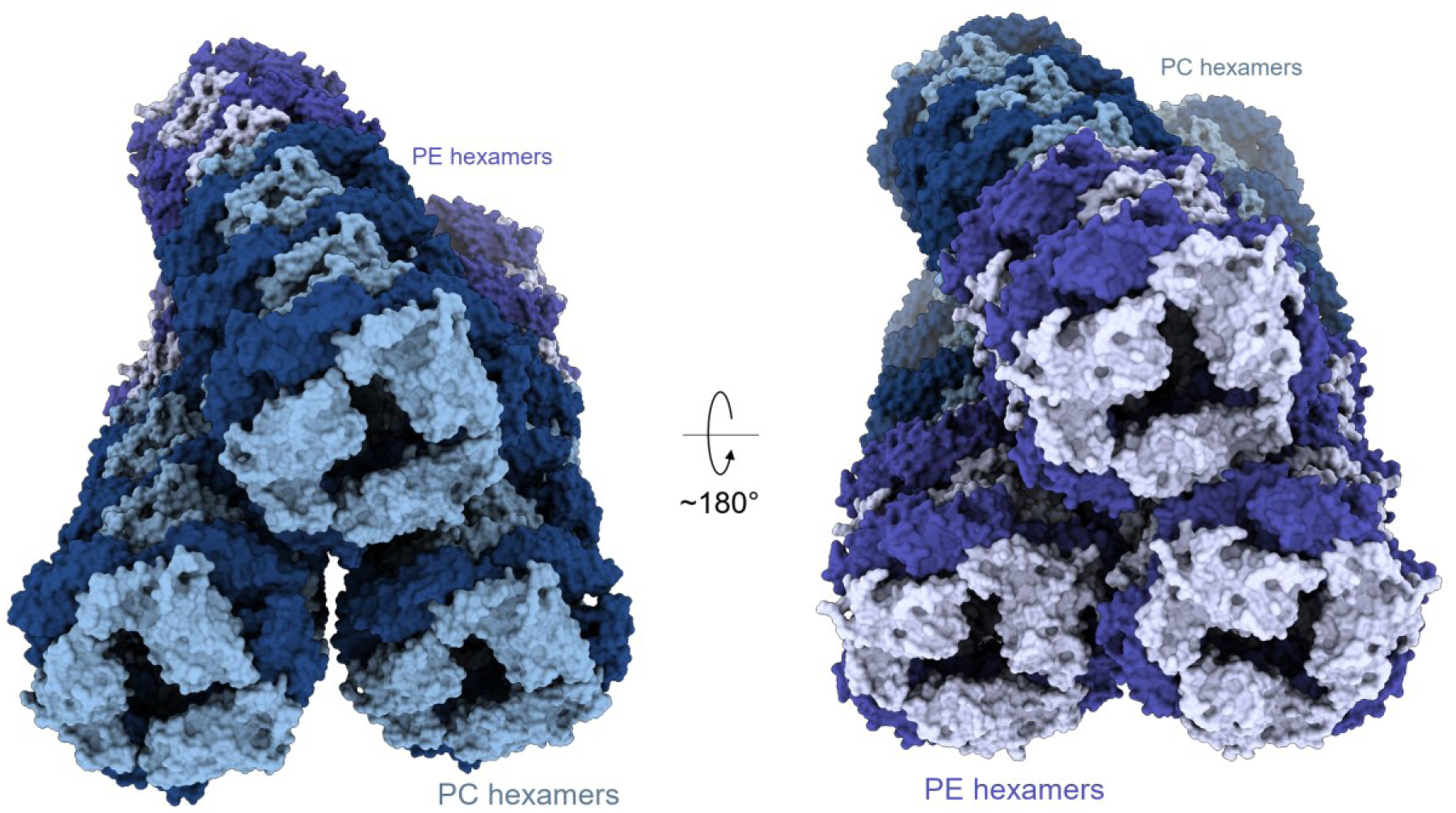
Helicity of rod bundles in GviPBS. Alpha and beta PBP subunits are shown by dark and light tints of color, respectively. Two views of a rod bundle showing non-parallel and helical orientation of the rods. The period of a superhelix is estimated as ∼1000 Å.

**Fig. S7.**
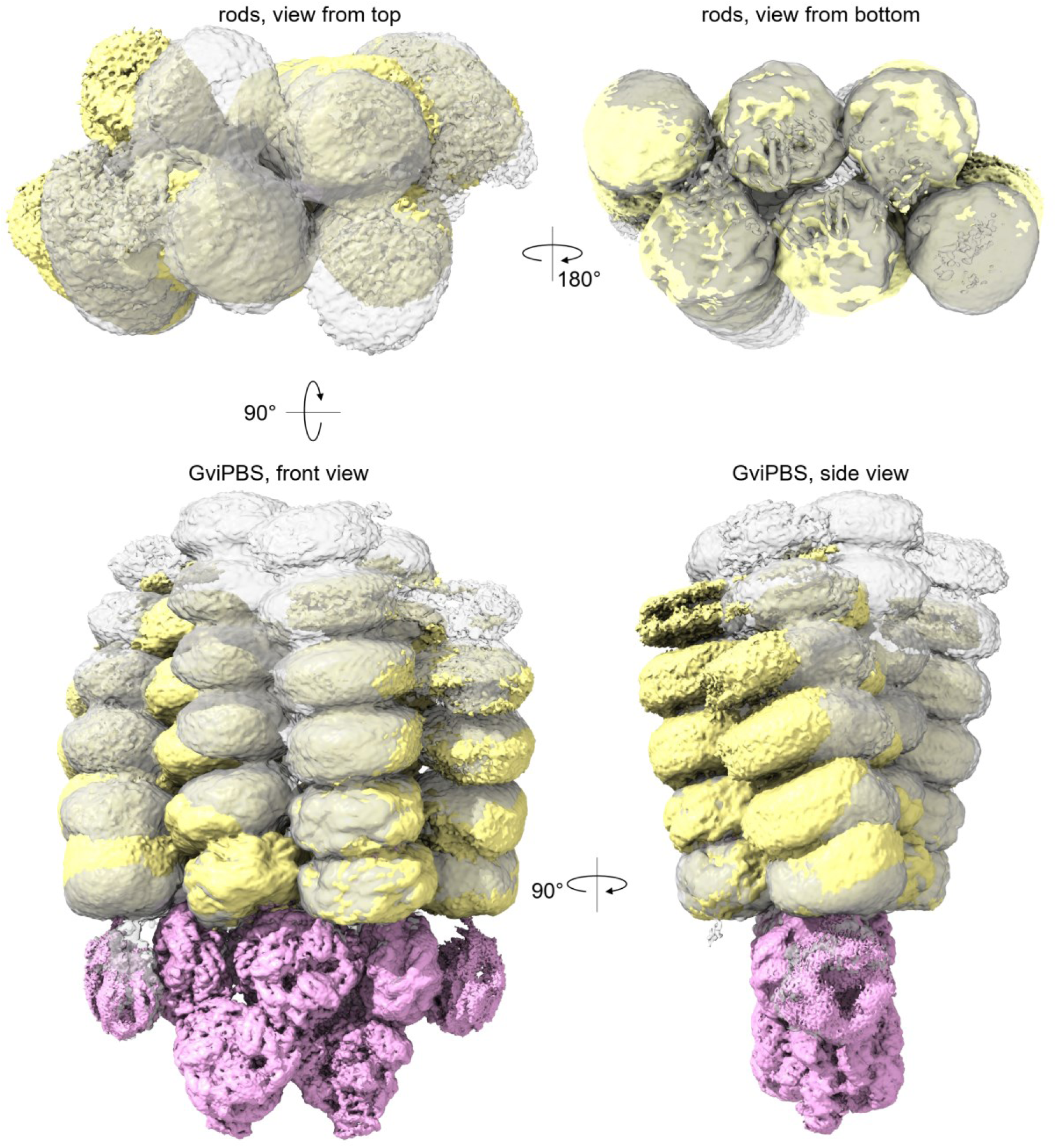
Conformational heterogeneity in rod bundles in GviPBS. Shown are different views of the superposition of two cryo-EM maps (light yellow and grey) corresponding to the two different conformational states of the rod bundles. Pink map corresponds to the AP core with Rx1/Rx1’ hexamers of PC; this map is omitted on the top views for clarity.

**Fig. S8.**
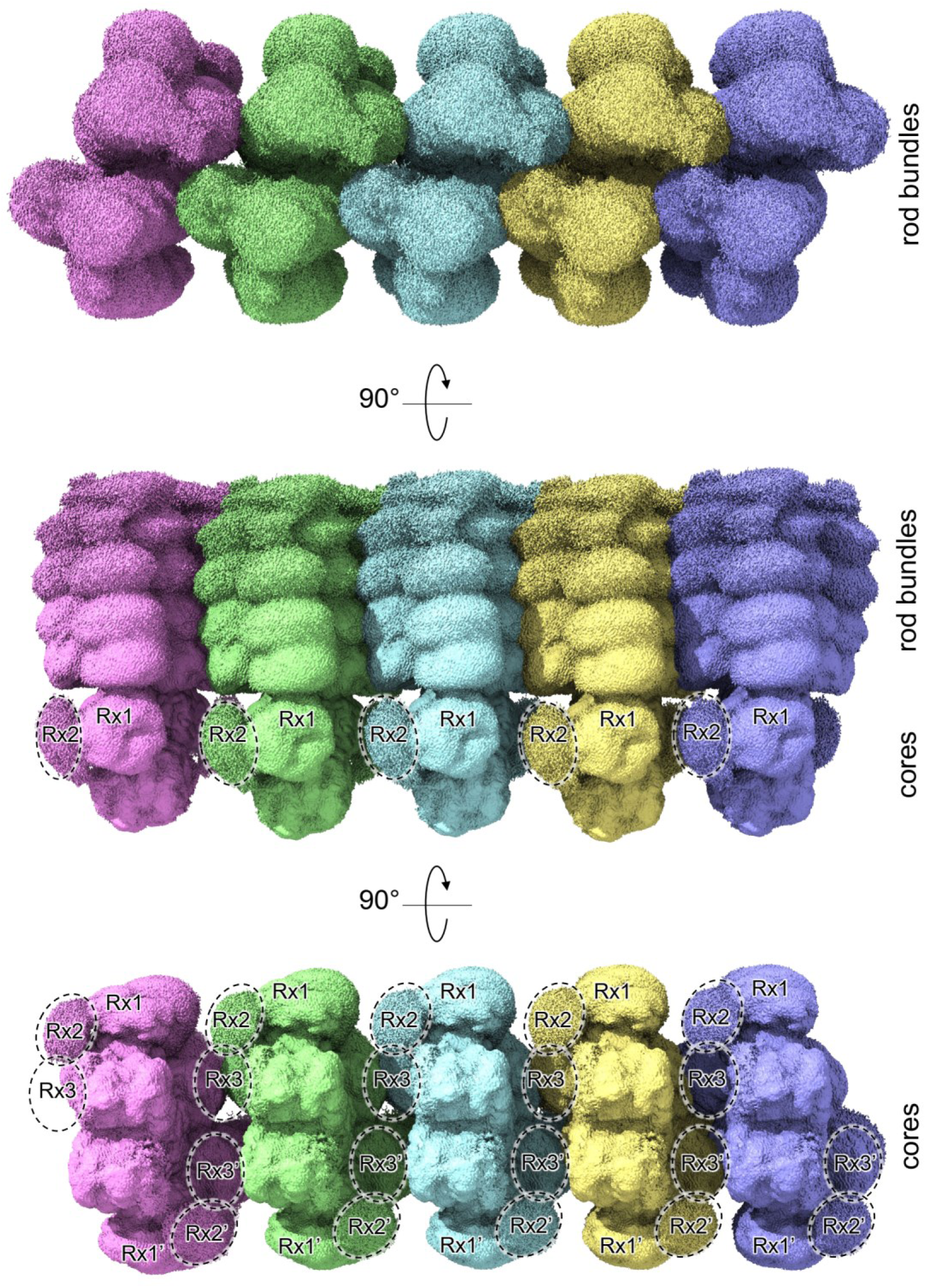
Tentative model showing the principle of GviPBS assembly into arrays. Five stacked copies of the GviPBS cryo-EM map are shown from different views. Note that the poorly resolved density in the vicinity of Rx1/Rx1’ suggests the position of the Rx2/Rx2’ hexamers. The third hexamers Rx3/Rx3’ are too dynamic and have no density, although their approximate location can be predicted from the presence of the third REP domain in the Glr2806 rod-core linkers (*16*). The complete GviPBS assemblies with six Rx hexamers (Rx1/Rx1’, Rx2/Rx2’, Rx3/Rx3’) belting the AP core can attach to each other in arrays, which would effectively use limited space in the plasma membrane in thylakoid-less *G. violaceus* cells, consistent with earlier considerations and observations (*1*, *12*).

**Fig. S9.**
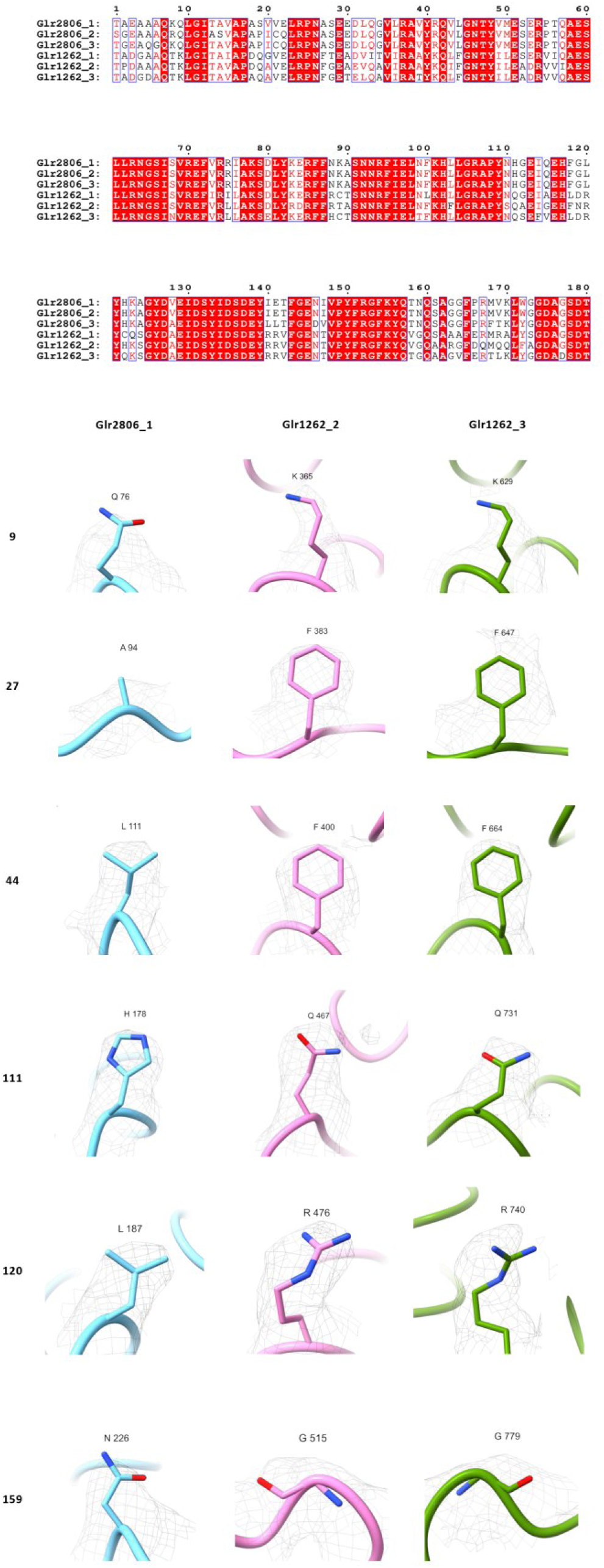
Identification of Glr1262 and Glr2806 linkers by cryo-EM density for the positions specifically conserved only within REP domains of either protein. Top, multiple sequence alignment of REP domains of the two proteins. Bottom, representative demarcating positions supported by cryo-EM density.

**Fig. S10.**
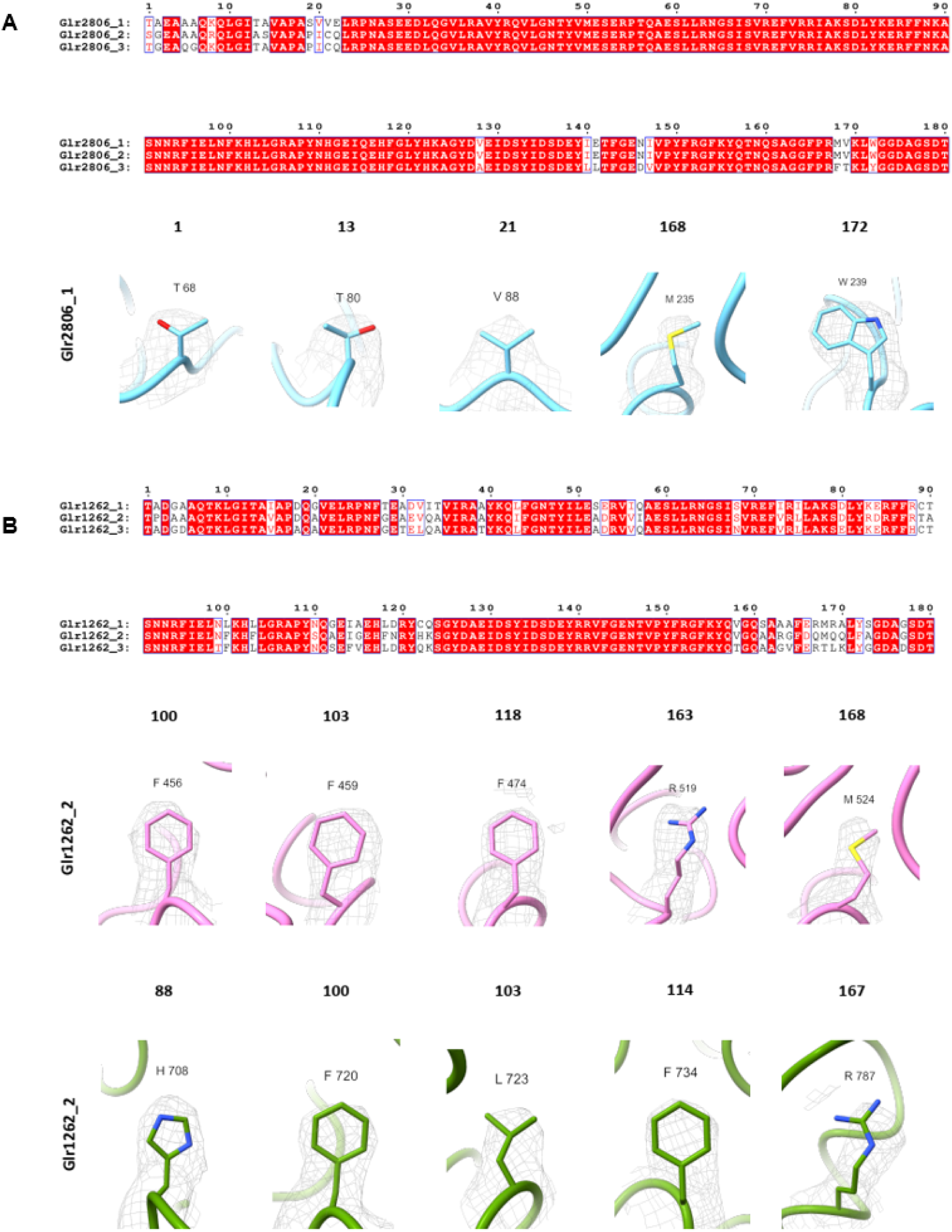
Identification of REP domains within either Glr1262 or Glr2806 linkers by cryo-EM density for the nonconserved positions. **A**. Top, multiple sequence alignment of REP domains of Glr2806. Bottom, representative demarcating positions supported by cryo-EM density. **B**. Top, multiple sequence alignment of REP domains of Glr1262. Bottom, representative demarcating positions supported by cryo-EM density.

**Fig. S11.**
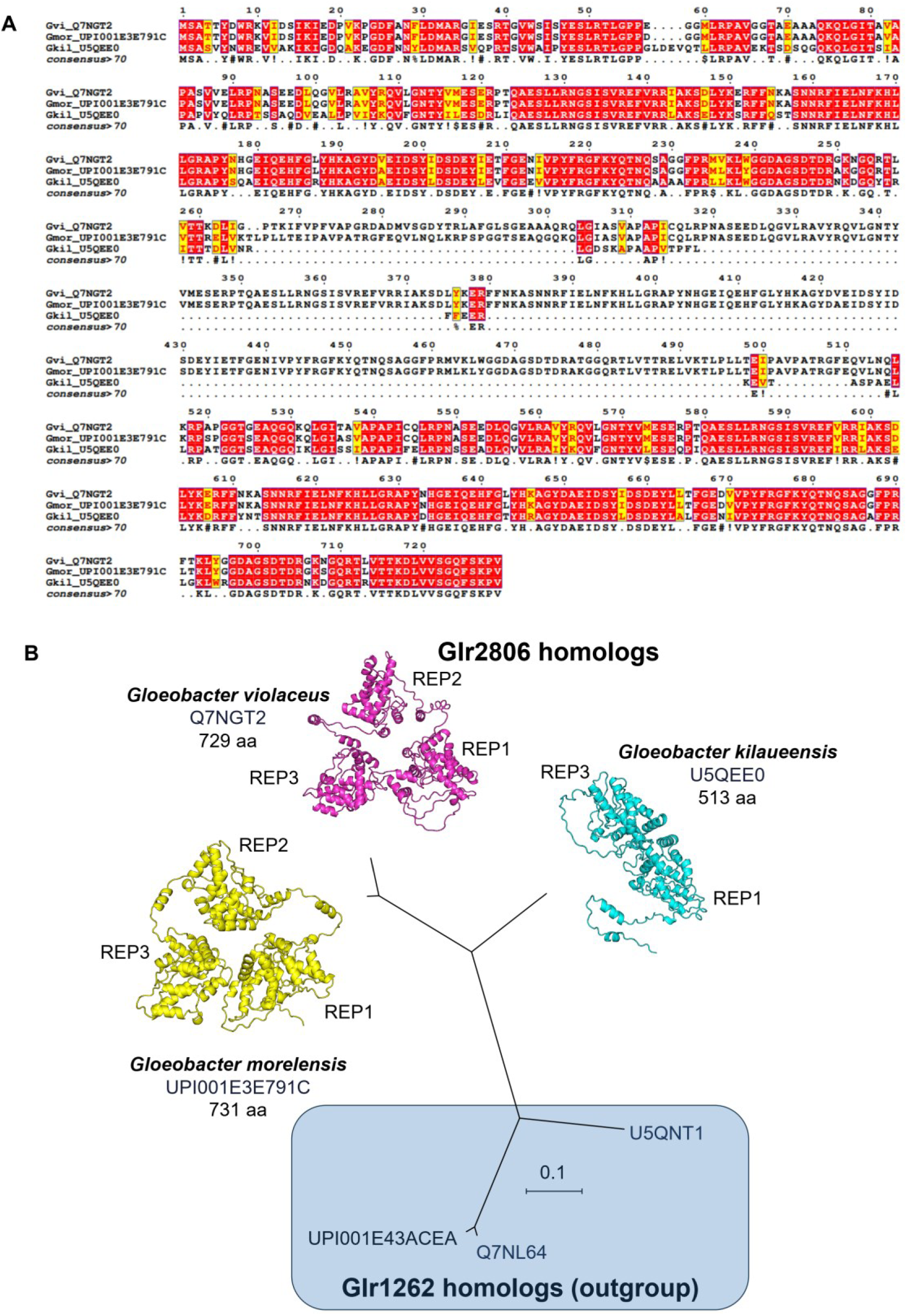
Analysis of the number of REP domains in Glr2806 linkers of *Gloeobacter* species. **A**. Multiple sequence alignment of the Glr2806 homologs from *G. violaceus*, *G. morelensis* and *G. kilaueensis* showing that the latter has only two REP domains out of three found in other two homologs. **B**. Phylogenetic tree of Glr2806 homologs built using Glr1262 homologs from the same organisms as an outgroup. Scale bar represents the number of per residue substitutions. Alphafold3 (*53*) models of the three Glr2806 homologs analyzed are shown along with the corresponding REP domains numbered.

**Fig. S12.**
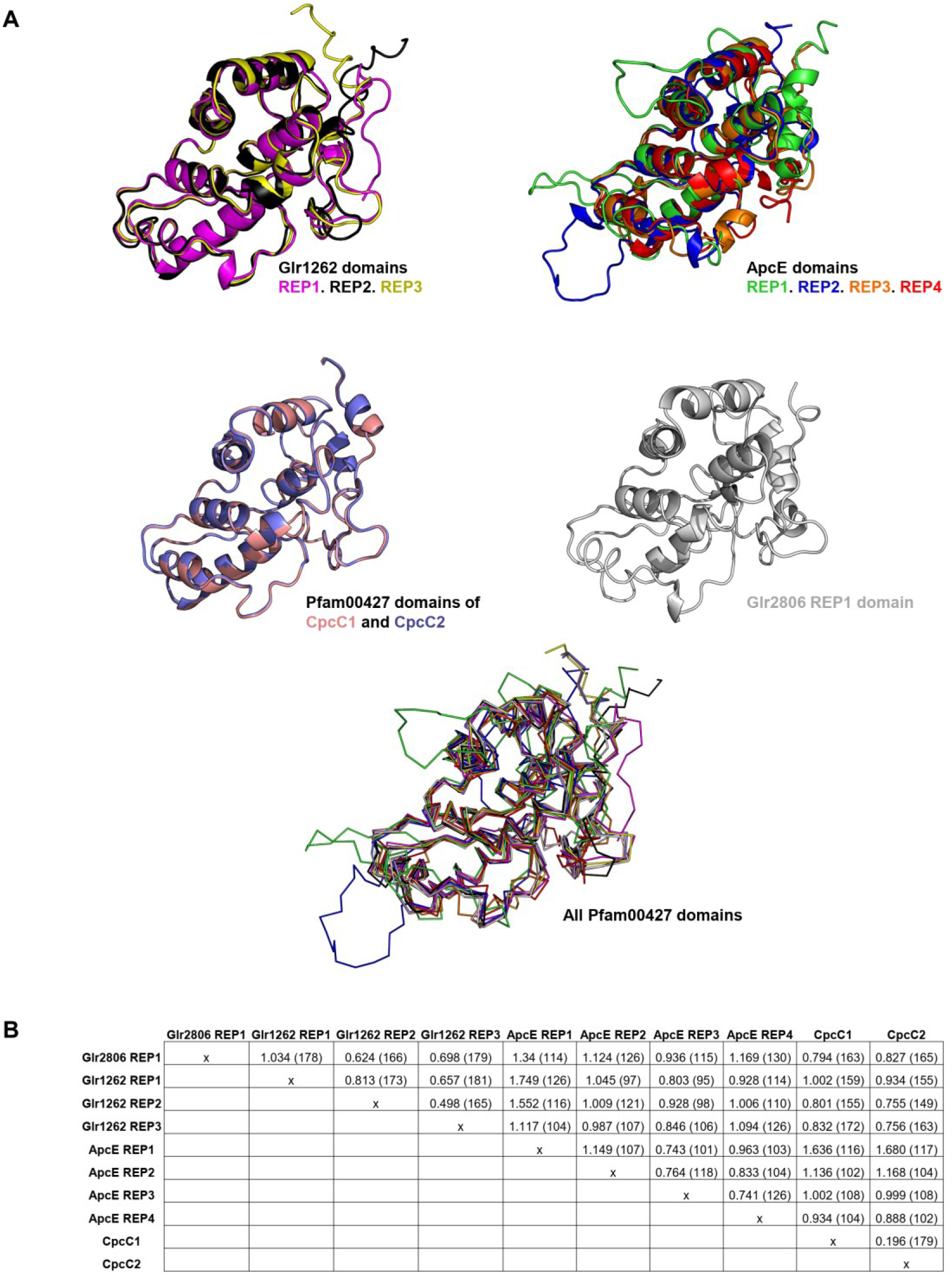
Pfam00427 domains of GviPBS linker proteins. **A**. Structural alignment of three REP domains of Glr1262, ApcE, CpcC1/CpcC2 and Glr2806 proteins. **B**. Cα RMSD values characterizing the similarity of the Pfam00427 domains found in GviPBS proteins. The values in Å were calculated upon structural alignment of a given domain pair. Numbers in parentheses indicate the number of the aligned atoms.

**Fig. S13.**
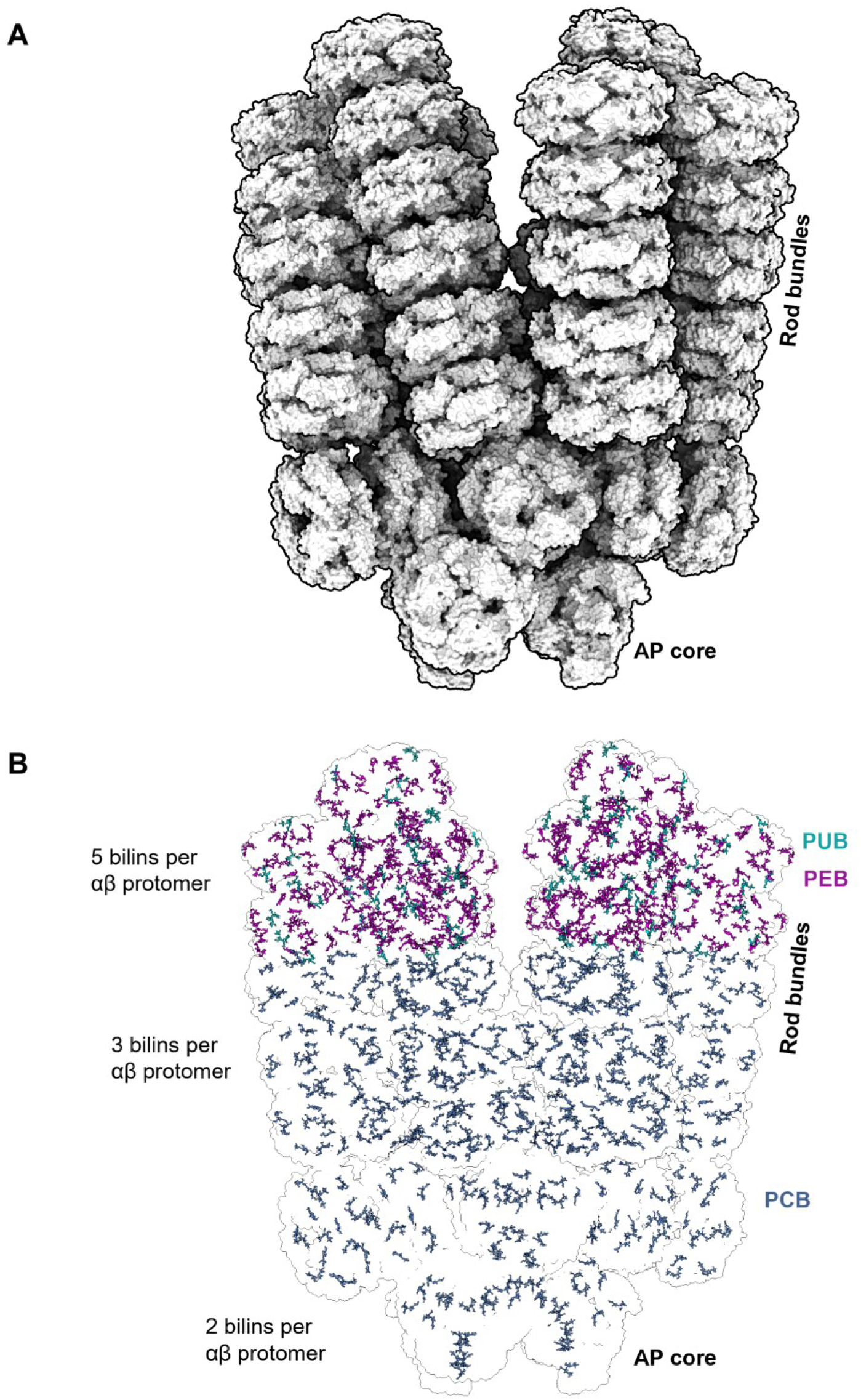
GviPBS bilins exhibit a descending concentration gradient from the rods to the core. **A**. GviPBS model in surface representation. **B**. Silhouette of the GviPBS model showing the location of PCB, PEB and PUB. Given the different numbers of bilins bound in PE, PC and AP, the gradient is formed.

**Fig. S14.**
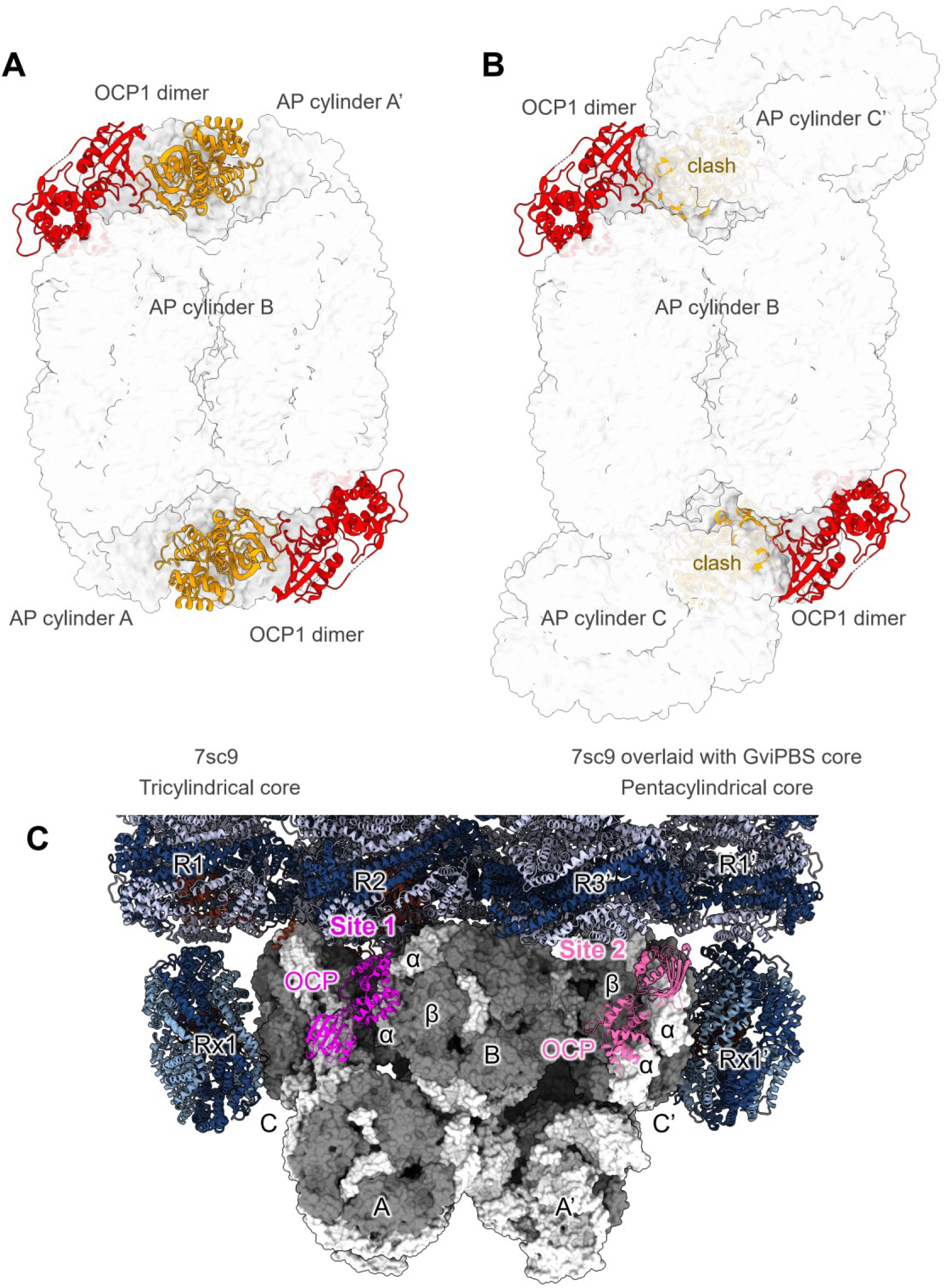
OCP-binding sites in tri- and pentacylindrical PBS cores. **A**. Cryo-EM structure of OCP1 bound to the tricylindrical core of SynPBS (PDB 7cs9 (*7*)). The AP cores are shown from the top, with the OCP dimer (subunits in red and yellow) bound to the top (B) and bottom (A/A’) cylinders. **B**. In the pentacylindrical PBS core, such as in GviPBS, the C/C’ cylinders of AP completely mask the OCP-binding site on the A/A’ cylinders and thereby block the attachment of the yellow OCP subunit, suggesting an existence of alternative OCP-binding sites. **C**. Structural alignment using conserved AP hexamers and the 7sc9 structure reveals a novel tentative OCP-binding site (Site 2) located on the side of the C/C’ cylinder, at the joint of two α and one β AP chains. This joint is equivalent to that of the OCP1-binding site on the top B cylinder observed in the 7sc9 structure (Site 1).

**Fig. S15.**
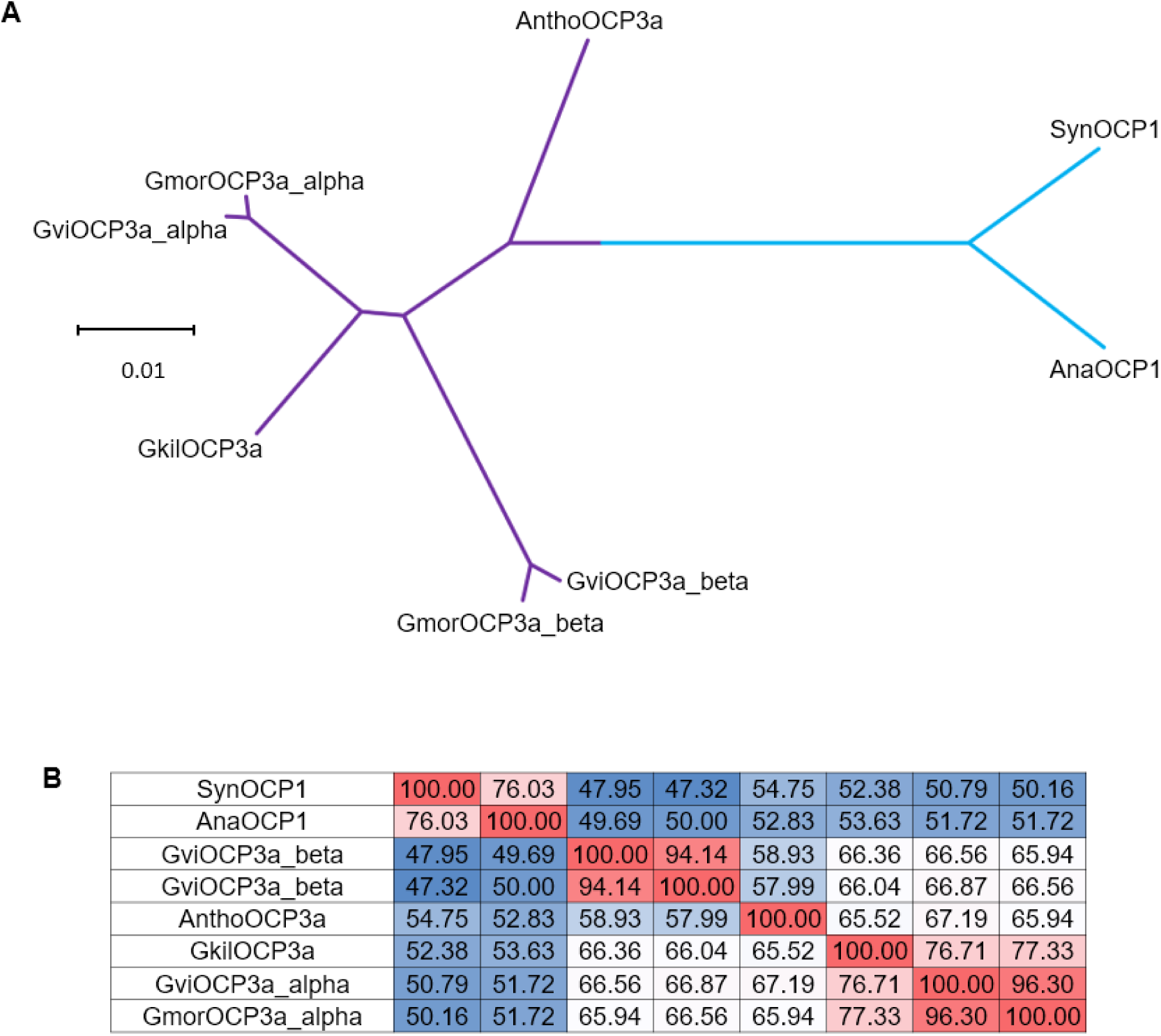
Phylogenetic relationships of the OCP3a variants. **A**. Phylogenetic tree showing distinct clusterization of OCP3a_α and OCP3a_β from different species of *Gloeobacter*, with two OCP1 representatives taken as an outgroup. **B**. Percent identity matrix of the analyzed sequences. SynOCP1 and AnaOCP1 are included for reference. Note that *G. kilaueensis* and *A. panamensis* (AnthoOCP3a) contain single OCP3a variants, of which the former most likely belongs to the α subclade, whereas the identity of AnthoOCP3a is less certain due to its intermediate position.

**Fig. S16.**
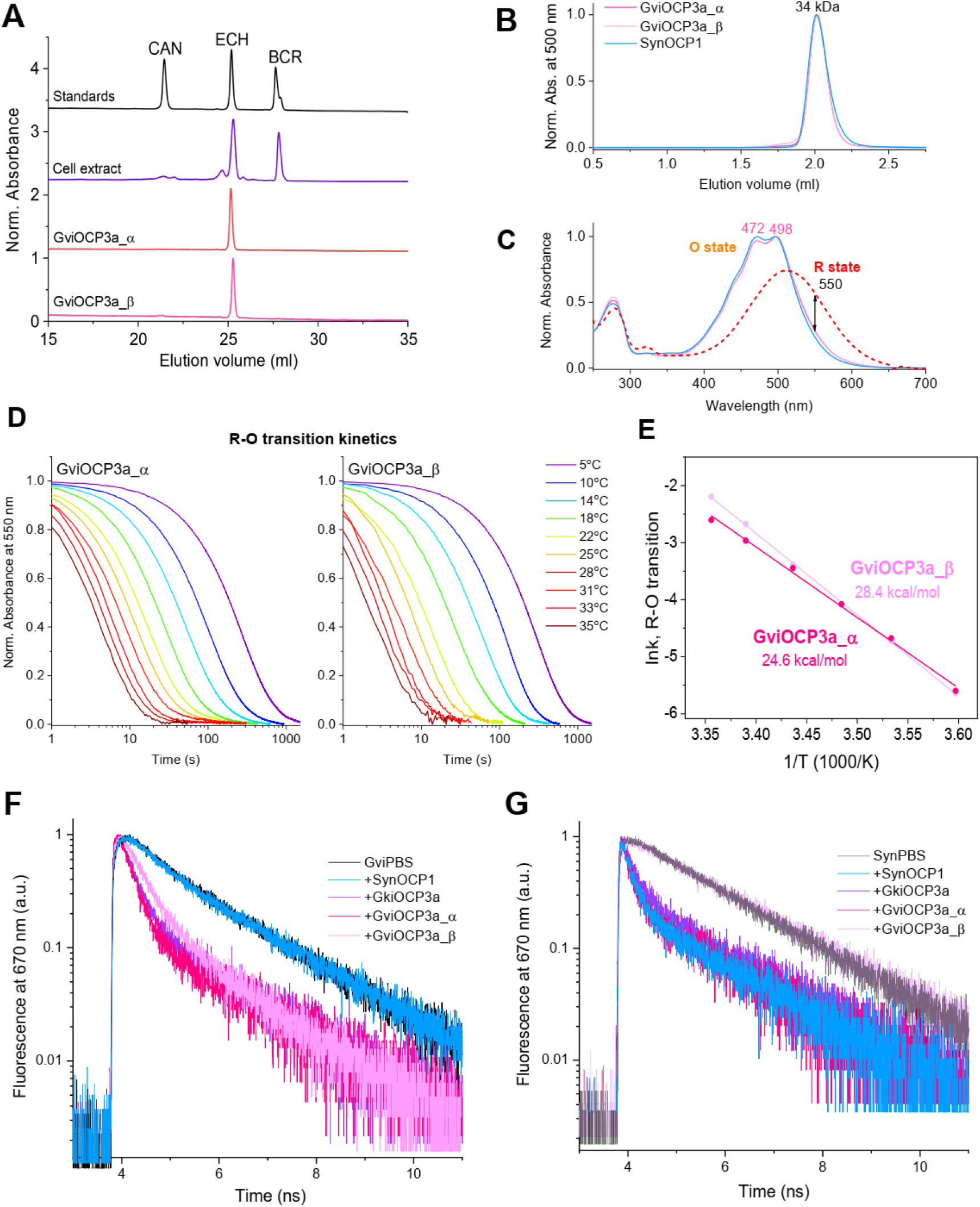
Characterization of GviOCP3a_α and GviOCP3a_β produced in carotenoid-synthesizing *E. coli* cells (*46*). **A**. HPLC profiles showing that both recombinant proteins bound exclusively echinenone (ECH) under the conditions used for protein expression. No canthaxanthin (CAN) binding was detected as this carotenoid did not have time to form from β-carotene (BCR) and echinenone (ECH) by the crtO ketolase in the expression system used (*35*, *46*). The elution profile for the carotenoid standards is shown. **B,C**. Analysis of recombinant OCP3a variants from *G. violaceus* (GviOCP3a_α and GviOCP3a_β) by size-exclusion chromatography (**B**) and absorbance spectroscopy (**C**. OCP1 from *Synechocystis* 6803 (SynOCP1) is shown for comparison. The apparent mass of OCP peak is indicated. The absorbance spectra of the dark-adapted orange and photoactivated red states of GviOCP3a_α are shown by solid (O state) and dashed lines (R state). Arrow indicates the wavelength used to monitor the R-O transition. **D**. Characterization of the photoactivity of the GviOCP3a variants by following the R-O relaxation after pre-adaptation to the actinic light (blue LED), at various temperatures as indicated. Note that the proteins exhibit similar photoactivity and relaxation rates in the whole range of temperatures tested. **E**. Arrhenius plot for GviOCP3a_α and GviOCP3a_β characterizing the temperature dependencies of their R-O transition. The activation energy barrier values are indicated. **F,G**. The ability of GviOCP3a variants to quench PBS from *G. violaceus* (**F**) or *Synechocystis* 6803 (**G**) studied by time-resolved fluorescence spectroscopy. The excitation wavelength was 570 nm. SynOCP1 and GkilOCP3a were used as controls. Note that while GviOCP3a_α is capable of quenching both PBS types, GviOCP3a_β is not capable of quenching SynPBS. GviOCP3a_β quenches GviPBS with an altered efficiency, suggesting that GviOCP3a_β binds and quenches GviPBS in an uncommon site.

**Fig. S17.**
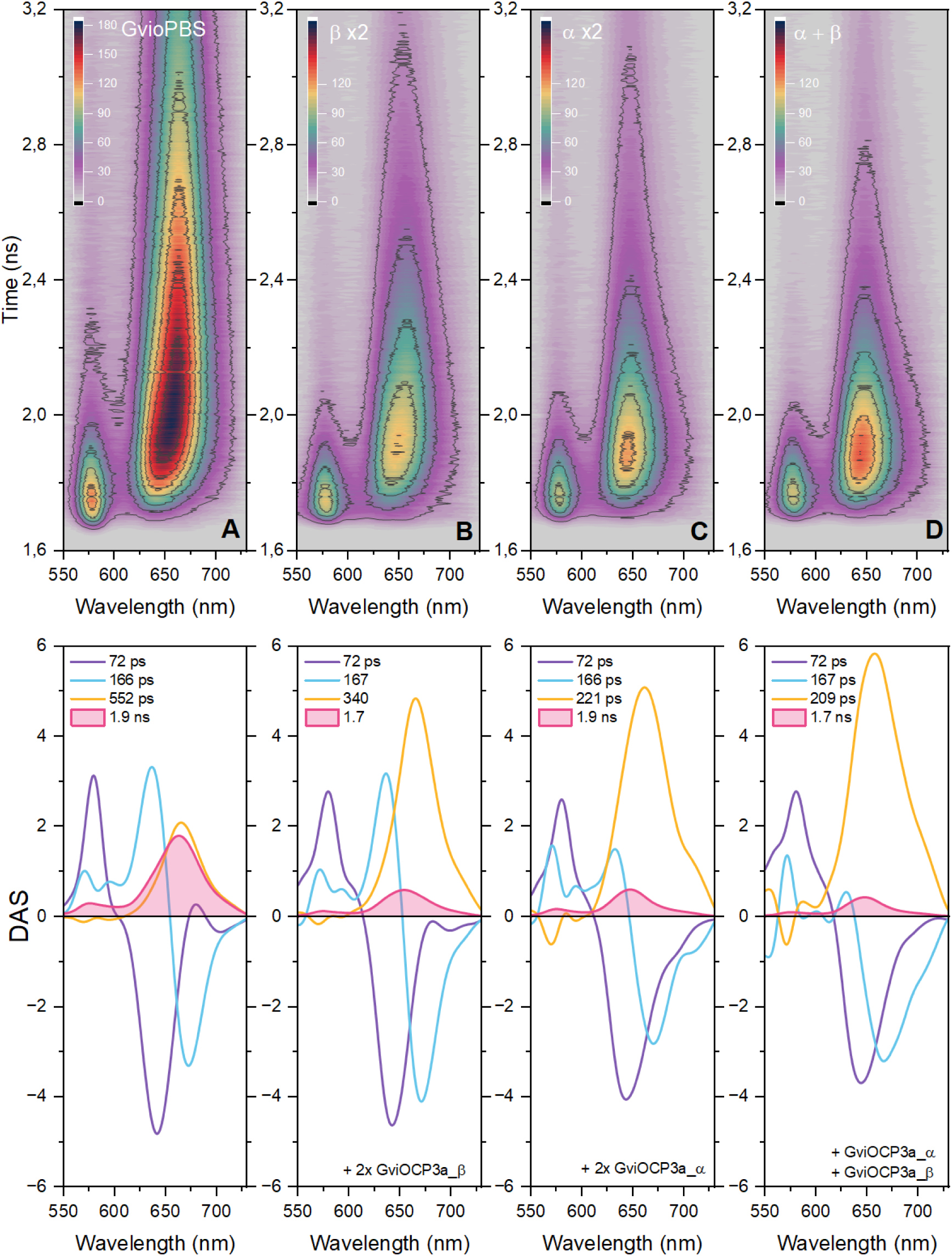
Time-resolved emission spectra (TRES) of GviPBS fluorescence in the absence (**A**), presence of the GviOCP3a_β (**B**), GviOCP3a_α (**C**), and their combination (**D**). Below each panel is a corresponding set of Decay associated spectra (DAS) of GviPBS fluorescence. The numbers indicate characteristic kinetic components derived from the global analysis of TRES showing the response to photoactivation of GviOCP3a_β, GviOCP3a_α and their combination. GviPBS fluorescence was excited at 485 nm by 3 mW 150 fs laser pulses at 80 MHz. The temperature of the samples was stabilized at 10 °C during all experiments to prevent fluorescence recovery.

**Fig. S18.**
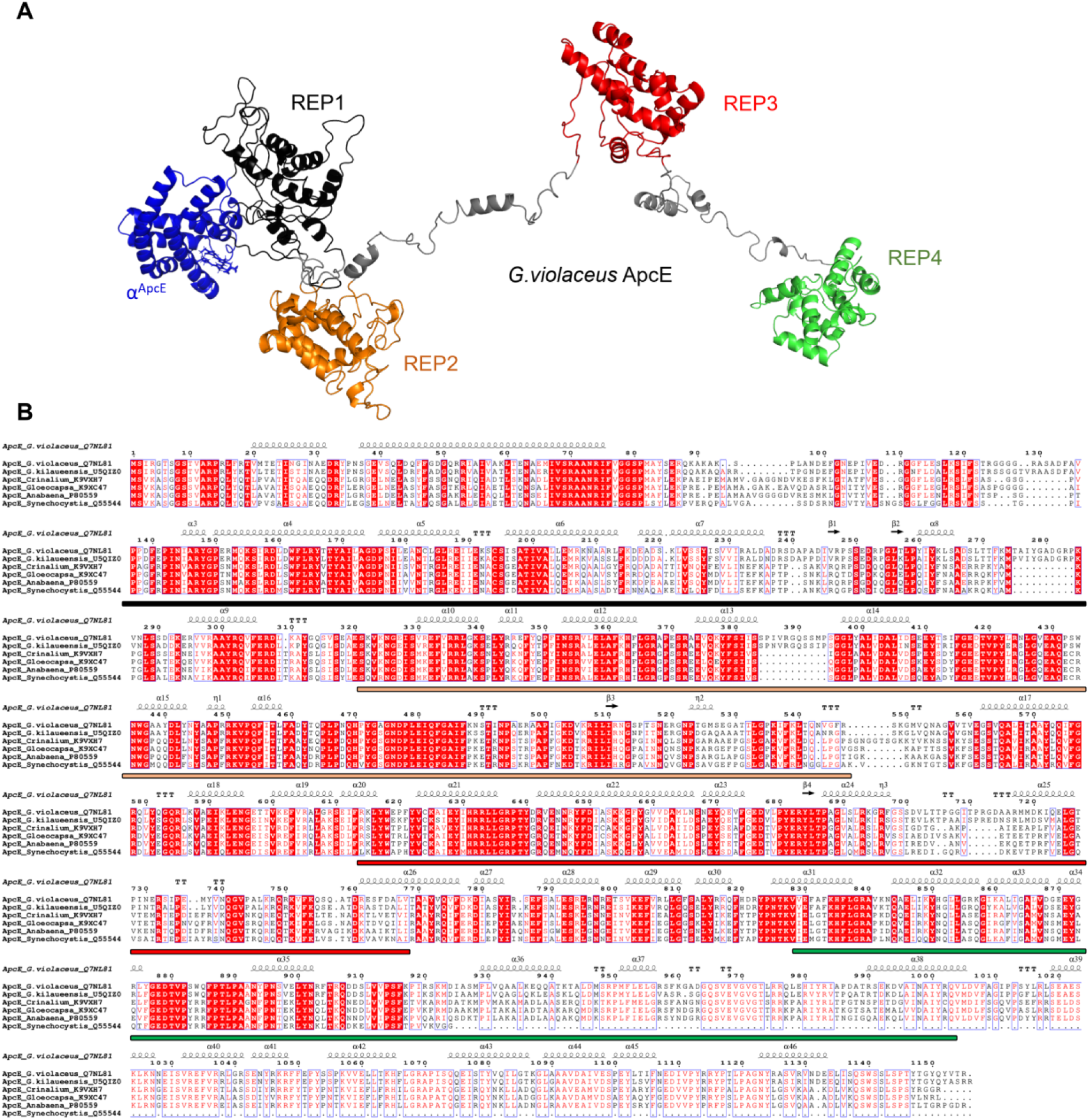
Domain structure of ApcE proteins. **A**. One copy of ApcE from *G. violaceus* showing the four REP domains following the N-terminal bilin-binding domain α^ApcE^. **B**. Multiple sequence alignment of ApcE linker proteins from species whose OCP proteins were studied in this work (*G. violaceus*, *G. kilaueensis*, *Crinalium epipsammum* PCC 9333, *Gloeocapsa* sp. PCC 7428, *Anabaena* 29413 and *Synechocystis* 6803). Given that the number of REP domains in ApcE proteins indicate the number of cylinders in the respective PBS core, all taken species possess the pentacylindrical PBS cores (four REP domains), except for *Synechocystis* 6803, which possesses the tricylindrical PBS core (the last REP domain is missing). The location of REP domains are indicated above the alignment with the color coding corresponding to panel A.

**Fig. S19.**
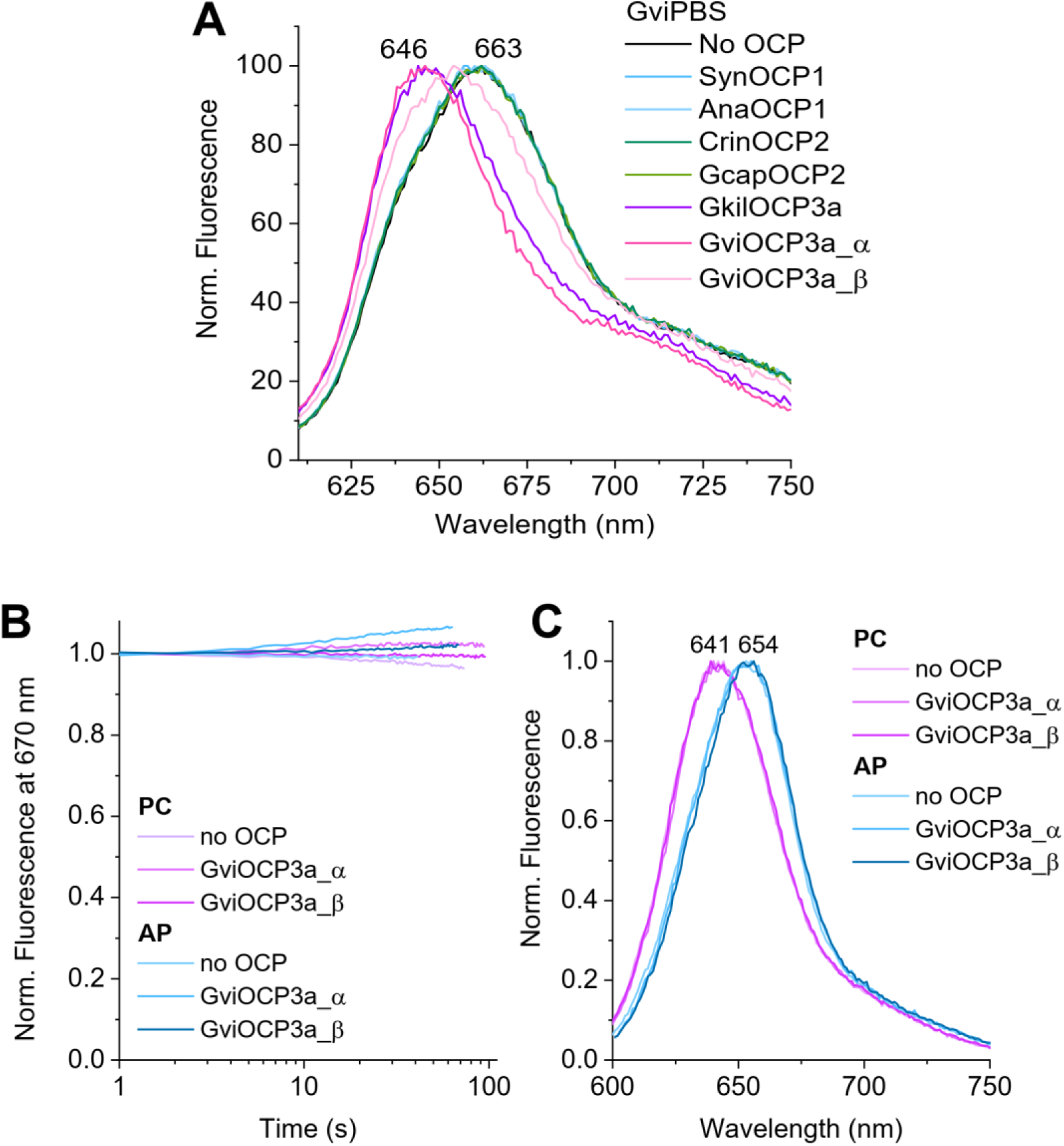
Effect of GviOCP3a_α and GviOCP3a_β on fluorescence of separated PC and AP components of GviPBS. Temperature 5 °C. **A**. Normalized fluorescence spectra of assembled GviPBS recorded in the absence or in the presence of either OCP variant, as indicated. Note that the maxima of quenched and unquenched GviPBS correspond to those of PC and AP components, respectively. **B**. Changes in the fluorescence intensity of the isolated PC or AP samples over time, with or without the addition of either GviOCP3a variant. **C**. Normalized fluorescence spectra of isolated PC and AP samples recorded in the absence or in the presence of either OCP variant, as indicated. Excitation in all experiments here was at 575 nm.

**Movie S1**. The cryo-EM reconstruction of GviPBS showing the overall complexity of its architecture, the precise location of the identified linker proteins in the rods and the core, as well as the conformational mobility of the rod bundles sampling conformational space from nearly parallel position to a position diverged by ∼15°.

